# Inhibitory and disinhibitory VIP IN-mediated circuits in neocortex

**DOI:** 10.1101/2025.02.26.640383

**Authors:** Shlomo Dellal, Hector Zurita, Ilya Kruglikov, Manuel Valero, Pablo Abad-Perez, Erez Geron, John Hongyu Meng, Alvar Prönneke, Jessica L. Hanson, Ema Mir, Marina Ongaro, Xiao-Jing Wang, György Buzsáki, Robert Machold, Bernardo Rudy

**Affiliations:** Neuroscience Institute, NYU Grossman School of Medicine, New York, NY, 10016; Hospital del Mar Medical Research Institute, Barcelona, Spain; Universidad Cardenal Herrera-CEU, CEU Universities, Spain; Center for Neural Science, NYU, New York, NY, 10003; Department of Neuroscience and Physiology, NYU Grossman School of Medicine, New York, NY, 10016; Department of Anesthesiology, Perioperative Care and Pain Medicine, NYU Grossman School of Medicine, New York, NY, 10016

**Keywords:** VIP interneurons, somatosensory cortex, cortical disinhibitory circuits, GABAergic microcircuits, interneuron diversity, intersectional genetic targeting, neocortical synaptic connectivity, neuromodulatory control, endocannabinoid signaling, optogenetic manipulation

## Abstract

Cortical GABAergic interneurons (INs) expressing the neuropeptide vasoactive-intestinal peptide (VIP) predominantly function by inhibiting dendritic-targeting somatostatin (SST) expressing INs, thereby disinhibiting pyramidal cells (PCs) and facilitating cortical circuit plasticity. VIP INs are a molecularly heterogeneous group, but the physiological significance of this diversity is unclear at present. Here, we have characterized the functional diversity of VIP INs in the primary somatosensory cortex (vS1) using intersectional genetic approaches. We found that VIP INs are comprised of four primary populations that exhibit different laminar distributions, axonal and dendritic arbors, intrinsic electrophysiological properties, and efferent connectivity. Furthermore, we observe that these populations are differentially activated by long-range inputs, and display distinct responses to neuromodulation by endocannabinoids, acetylcholine and noradrenaline. Stimulation of VIP IN subpopulations *in vivo* results in differential effects on the cortical network, thus providing evidence for specialized modes of VIP IN-mediated regulation of PC activity during cortical information processing.

## Introduction

The remarkable diversity of GABAergic inhibitory interneurons (INs) in the cerebral cortex has been appreciated for over a century ^1,2^. Cortical INs are comprised of distinct populations that provide tailored inhibition to pyramidal cells (PCs) and other INs, thereby enabling precise control of cortical circuit activity. The specialization of INs with regard to dendritic and axonal morphology, compartment-specific connectivity, and intrinsic electrophysiological and synaptic properties allows for the generation of an impressive variety of circuit motifs and is thereby thought to contribute to the enormous computational power of the neocortex ^3,4^.

Recently, among the major types of INs in neocortex, the INs that express the neuropeptide vasoactive intestinal peptide (VIP) have attracted attention following the discovery that they mediate a disinhibitory circuit in multiple cortical areas. As a group, VIP INs do not provide substantial inhibition onto pyramidal cells (PCs), but instead preferentially inhibit INs expressing the neuropeptide somatostatin (SST INs), a key source of dendritic inhibition in the cortex. Several studies published in 2013-2015 reported that VIP INs in the neocortex are major targets of feedback connections and behavioral state-dependent neuromodulator release, and that their inhibition of SST INs and consequent disinhibition of PCs is a key mechanism of top-down and context-dependent processing ^5–11^. The studies were performed in L2/3, where VIP INs are enriched and are the largest IN population across different cortical areas (somatosensory barrel cortex, vS1; primary visual cortex, V1; medial prefrontal cortex, mPFC; and primary auditory cortex, A1). This work confirmed conclusions from earlier anatomical studies performed in the hippocampus ^12–17^ that showed that INs expressing VIP preferentially innervated other GABAergic neurons, and suggested important roles for this disinhibitory circuit mechanism in the integration of sensory and motor signals in vS1, behavioral state-dependent control of visual responses ^18^ and synaptic plasticity in V1^19,20^, visual top-down attention, and in the regulation of cortical activity by reinforcement signals ^21^, respectively. Since then, numerous publications have appeared documenting the existence and functional significance of this canonical VIP-SST disinhibitory circuit, expanding the brain areas and behaviors where this motif is thought to be important. For instance, recent studies have found that VIP-dependent disinhibition is important in gating associative synaptic plasticity in olfactory cortex ^22^, in goal-oriented spatial learning in the hippocampus ^23^, in mediating the effects of neuropeptides in the associative learning of fear memories in the auditory cortex ^24^, in gating the ability of hippocampal input to generate prefrontal representations which drive avoidance behavior ^25^ and in sensory processing during adaptive behavior in the anterior insular cortex ^26^. Underscoring their importance in cognition, VIP IN dysfunction has been implicated in schizophrenia ^21,27^, Rett syndrome ^28^, and in the cognitive effects of Dravet Syndrome ^29,30^.

Most of these studies have treated VIP INs as a homogeneous neuronal population and relied on manipulations of total VIP INs in VIP-Cre mice. However, there is compelling evidence that VIP INs are diverse. Immunohistochemical anatomical studies in the Freund laboratory ^12,31^ provided evidence for the existence of two major types of VIP INs in the hippocampus, with the majority of the VIP cells co-expressing the Ca2+ binding protein calretinin (CR or Calb2) and a much smaller group co-expressing the neuropeptide CCK. Intersectional genetics based on the co-expression of VIP and CR or CCK showed that these markers also defined distinct VIP INs in the neocortex, where the VIP/CCK group is more abundant than in the hippocampus ^32^. More recently, single cell transcriptomic analyses have provided evidence for extensive molecular heterogeneity of VIP INs in the neocortex ^33–36^. However, the relationship between the VIP-IN molecularly defined groups and functionally-relevant categories remains to be studied and thus the physiological relevance of VIP IN diversity is not well understood at present.

Here we leveraged the accumulated transcriptomic insights from the Allen Institute with intersectional genetics to investigate the diversity of VIP INs in the neocortex. We found that VIP INs can be divided into four main populations: a group that expresses CR, two distinct groups that express CCK, and a group that does not express CR or CCK, but expresses the genes Cxcl14 and CRH. VIP neurons in each group differ in laminar distribution, morphological and electrophysiological properties and exhibit distinct connectivity patterns. VIP/CR INs almost exclusively target SST INs, whereas the non-CCK non-CR VIP INs also mainly target SST INs but also have connections to parvalbumin (PV) expressing INs. These two groups have essentially no connectivity to pyramidal cells (PCs). On the other hand, the two types of VIP/CCK INs target PCs as much or more than SST INs, but differ in their synaptic properties. Notably, we find that these VIP IN populations appear to target different SST IN subtypes, indicating the existence of several parallel VIP-SST disinhibitory circuits. Our studies also provide evidence that VIP IN subtypes are differentially driven by long range cortical inputs, as well as by neuromodulators such as acetylcholine, noradrenaline and endocannabinoids. Consistent with their specialized input and output properties, activation of distinct VIP IN subpopulations *in vivo* produced distinct effects on the cortical network. Together, our findings demonstrate the existence of multiple inhibitory and disinhibitory VIP IN-mediated circuits in the neocortex, and reveal an unexpected substructure in the regulation of information flow in the cortical network by VIP INs.

## Results

### Heterogeneity of VIP INs in the neocortex

VIP INs account for ∼12% of the GABAergic neurons in the neocortex, and are highly enriched in superficial cortical layers, with ∼60% present in L2/3, where they are the most abundant IN population (50% of all L2/3 INs) ^3^. L4, particularly its superficial part, is also enriched in VIP INs ^3,37^. In this study, in addition to L2/3, which has been the focus of most VIP IN studies in neocortex, we also included VIP cells in L4, which, as shown below resemble those in deep L2/3. VIP cells in L2-4 account for ∼80% of VIP INs in the neocortex and comprise the vast majority of the VIP cells that regulate the activity of L2/3 associative circuits. The sparse deeper layer (L5-6) VIP INs largely do not project to L2/3 and thus participate in distinct circuits ^37^.

Studies in the neocortex and the hippocampus have shown that VIP INs can be divided in two populations that differ in morphological and intrinsic electrophysiological properties based on co-expression of the neuropeptide CCK or the Ca2+-binding protein calretinin (CR) ^12,31,32^. Here, we first asked whether VIP/CCK and VIP/CR neurons account for all the VIP INs in L1-4 of the mouse barrel cortex (vS1; S1BF). Comparison of the cells labeled in VIP-Flp; CR-Cre; Ai65 and VIP-Flp; CCK-Cre; Ai65 mice to the total VIP population (labeled in VIP-Cre; Ai9 or VIP-Flp; Ai65F animals) in L2-4 of vS1 suggested there were VIP INs that were neither CR- or CCK-expressing, particularly in the middle part of L2/3. To investigate this possibility, we crossed the VIP-Flp; CR-Cre and the VIP-Flp; CCK-Cre mice to the intersectional/subtractive dual reporter FTLG ^38^ (Figure 1A). With this reporter, the intersected population (containing both Cre and Flp, e.g., the VIP/CR or the VIP/CCK cells in the VIP-Flp; CR-Cre or the VIP-Flp; CCK-Cre mice, respectively) is labeled with GFP, while the remaining cells that express Flp but not Cre are labeled with tdTomato. We next generated mice with both CR-Cre and CCK-Cre paired with VIP-Flp (VIP-Flp; CCK-Cre; CR-Cre), and crossed these animals with the FLTG reporter. This led to the discovery of a sizable population (∼30%) of VIP cells in L2-4 that does not express either CR or CCK (the non-CCK non-CR VIP neurons; Figure 1B). Implementing these labeling approaches allowed us to visualize, characterize and quantify the proportions of all three VIP subpopulations across the vS1 (S1BF) cortex (Figure 1C-F). Interestingly, the proportions of this non-CCK non-CR VIP population varied across different cortical areas, comprising ∼20% of total VIP INs in primary visual cortex but only ∼ 8% in prelimbic cortex. Work on transcriptomic differences across VIP subpopulations has revealed extensive molecular heterogeneity ^34,35,39^, including between the genetically defined VIP/CCK and VIP/CR groups ^36^. Examining the categorical frameworks established by the Allen Institute ^34,35^ and gene expression levels for CCK and Calb2, we generated an approximate correspondence between the latter and the VIP subpopulations identified here through cumulative intersectional genetics (Supplementary Figure 1).

**Figure 1.**
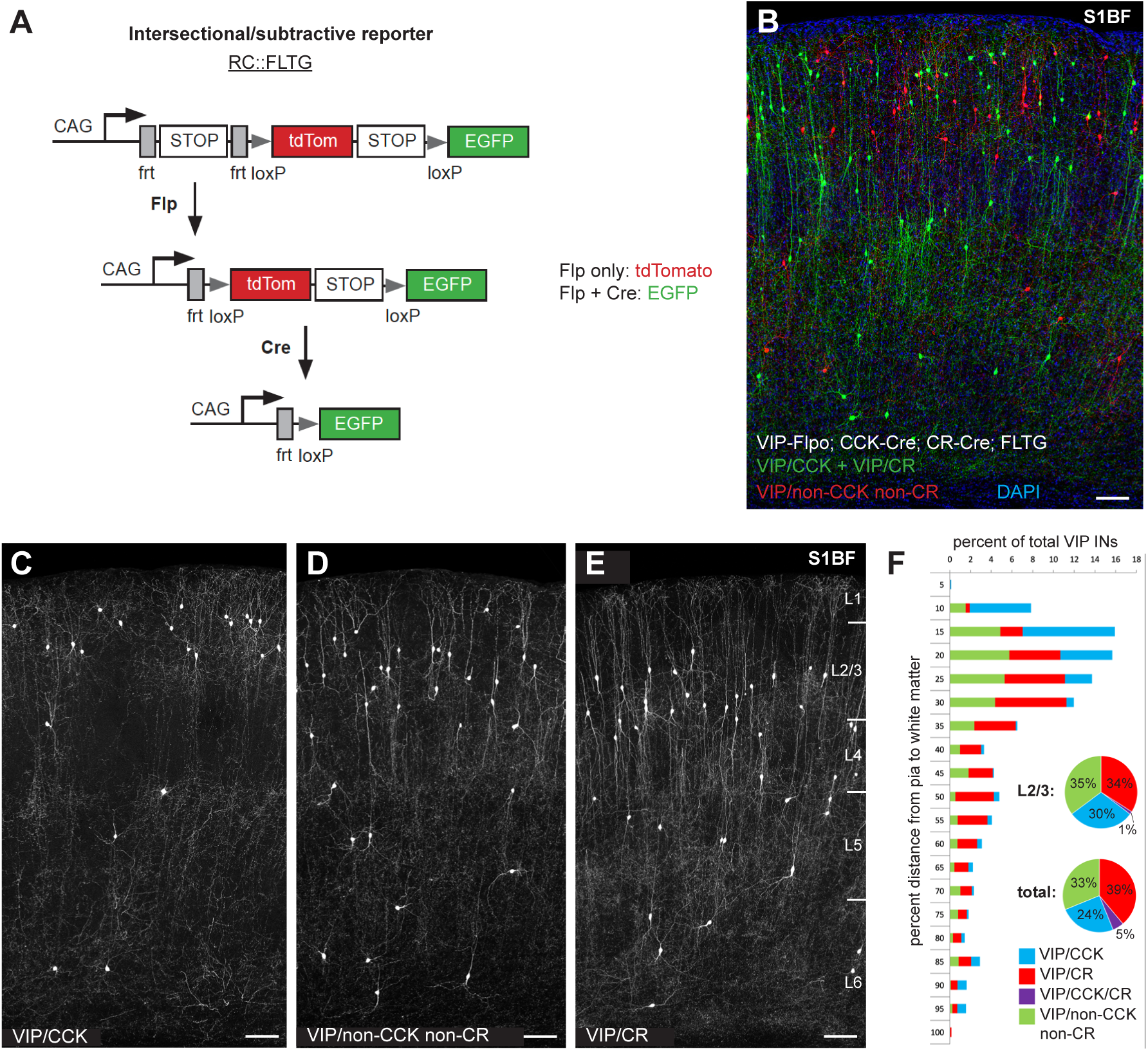
Resolution of distinct VIP IN populations using intersectional genetics. **(A)** Schematic of the FLTG reporter. Flp activity results in tdTomato expression and Flp + Cre results in EGFP expression. **(B)** Image of a brain tissue section (vS1) from VIP-Flpo; CCK-Cre; CR-Cre; FLTG animals illustrates the VIP/non-CCK non-CR population (red). **(C-E)** Images of VIP/CCK **(C)**, VIP/non-CCK non-CR **(D)** and VIP/CR **(E)** INs labeled using the FLTG reporter (50 μm thick sections of vS1 adult cortex). **(F)** Graphical representation of the proportions of each VIP subtype shown in C-E from the pia to white matter in 5% increments. Pie charts summarize the overall proportions of each subtype in L2/3 and overall (vS1), and include the inferred VIP/CCK/CR overlap. Scale bars in B-E represent 100 μm.

### Differential laminar distribution and morphological properties of VIP IN populations

The three genetically defined populations of VIP INs described above differ in their distribution within L2-4 of vS1 (Figure 1 C-F). VIP/CCK cells are most superficial, and are present in superficial L2, often near the border with L1; VIP/CR are present mainly in lower layer 2/3 and L4, and the non-CCK non-CR VIP neurons tend to occupy the middle part of L2/3. We performed extensive reconstructions of individual VIP INs from each of the three crosses, and found that these genetically-defined VIP IN subtypes exhibited notable differences in the morphologies of their dendritic and axonal arbors (Figure 2), predicting subtype specificity in input and output connectivity. Most VIP INs in L2-4 have a bipolar/bitufted dendritic morphology, with 2-3 vertically oriented primary dendrites (Figure 2; Supplementary Figures 2-4). The dendritic arbor of these neurons tends to be narrow and cross several layers in either direction (Figure 2). Their ascending dendritic branch extends fully through L1, often reaching close to the pial surface. In L1, or slightly before that, the dendrite branches extensively forming a dendritic tuft that could be targeted by the many cortico-cortical and subcortical projections to this layer ^40^. However, some VIP/CCK cells (about a third of the sampled neurons in this group) have a multipolar dendritic morphology with dendrites extending in all directions with a bias towards pia (Figure 2A; Supplementary Figures 2B-C, 3F and 4). Similarly, most of the L2-L4 VIP cells have a descending, vertically oriented and narrow axonal arbor, with branches that go deep in the column, predicting a direct influence which is vertically broad but laterally restricted (Figure 2A-C; Supplementary Figures 2 C-E, 3 G-I, 4). However, the axon of the multipolar VIP/CCK cells is local, largely restricted to L2/3 and is horizontally oriented, (with only a few collaterals going to deeper layers in some of the cells (Figure 2 A; Supplementary Figures 2 A-B and 3 F)).

**Figure 2.**
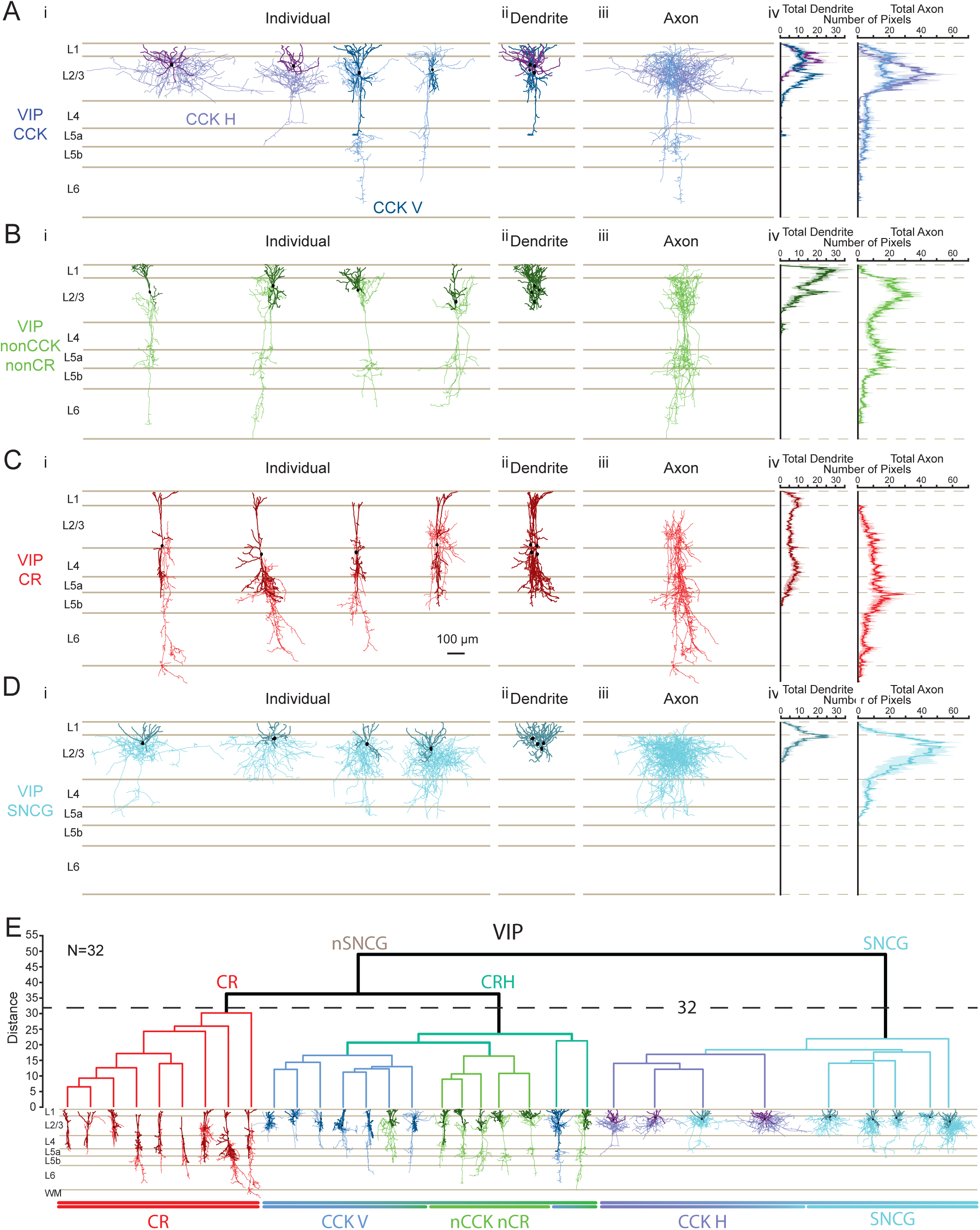
Morphological Properties of VIP IN populations. **(A)** VIP/CCK-expressing neurons. **(i)** Complete Neurolucida reconstructions of four CCK-expressing neurons reconstructed from VIP-Flp/CCK-Cre/FLTG mice across cortical layers. CCK+ neurons were classified as CCK Horizontal (CCK H) and CCK Vertical (CCK V), with their respective dendrites (dark lavender for CCK H, dark blue for CCK V), axons (light lavender for CCK H, light blue for CCK V), and soma (black) **(ii)** Superimposed dendrites and somata of the four neurons. **(iii)** Superimposed axons of the four neurons Quantification of pixel distribution for total reconstructed dendrites (left) and axons (right) in **(iv)** is shown for CCK H and CCK V neurons. **(B)** Same as **(A)** for non-CCK, non-CR VIP neurons reconstructed from VIP-Flp/CCKCR-Cre/FLTG mice, with dendrites shown in dark green, axons in light green, and soma in black. **(C)** Same as **(A)** for CR-expressing neurons reconstructed from VIP-Flp/CR-Cre/FLTG mice with dendrites shown in dark red, axons shown in light red, and soma in black. **(D)** Same as **(A)** for four SNCG-expressing neurons reconstructed from VIP-Cre/SNCG-Flp mice with dendrites shown in dark cyan, axons shown in cyan, and soma in black. **(E)** Unbiased hierarchical clustering (Ward’s method) was performed on all reconstructed neurons (n = 32) using a comprehensive set of morphological parameters, including somatic localization, surface area, volume, number of primary dendrites, total dendritic length, mean dendritic length, dendritic surface area, dendritic volume, dendritic polarity, total axonal length, axonal surface area, axonal volume, and axonal polarity. The Gap Statistic was used to determine the optimal number of clusters, leading to the selection of a distance threshold of 32 for final group separation. The most pronounced morphological distinction was observed between VIP/SNCG-like and VIP/non SNCG neurons, followed by the separation of VIP/CR and VIP/CRH-like neuronal morphologies. Below the clustering dendrogram, each corresponding morphology is displayed for visualization.

The finding of two different morphologies in the VIP/CCK group (neurons with multipolar dendritic harbor and horizontally oriented local axonal arbor (VIP/CCK H) versus neurons with a vertically oriented bipolar dendritic harbor and vertically oriented descending axonal arbor (VIP/CCK V) raised the possibility that VIP/CCK neurons consists of two distinct populations. Interestingly, transcriptomic analysis also supports the existence of two groups of VIP/CCK INs (Supplemental Figure 1). In addition to the IN clusters in the VIP subclass of GABAergic neurons, cells strongly expressing both VIP and CCK are found in the Synuclein γ (SNCG) IN subclass ^34,35,39^, an IN group that contains CCK basket cells ^41^. To explore whether the morphological differentiation we saw within the VIP/CCK group is related to this transcriptomic differentiation, we generated VIP-Cre; Sncg-Flpo mice, crossed these with the intersectional tdTomato reporter Ai65 and characterized the morphological properties of tdTomato labeled neurons. VIP/SNCG neurons were localized in superficial layer 2, near the border with L1 (Supplementary Figure 2 F-G), as observed for some of the INs labeled in the VIP/CCK intersectional mice, and resembled morphologically the multipolar VIP/CCK cells with a multipolar dendrite and horizontally oriented dendritic and axonal arbors (Figure 2 A & D; Supplementary Figure 2 A-B). These morphological features suggest that the horizontal VIP/CCK cells likely correspond to a set of VIP- and CCK-expressing “small basket cells” ^42–46^.

While the differences in dendritic and axonal arbor between the various VIP bipolar/bitufted cells are more subtle than their differences with the multipolar VIP/CCK neurons, they might be physiologically significant. The descending dendritic process of VIP/CR neurons is longer, often reaching deeper in the column, while it is restricted mostly to L1-3 in the bipolar VIP/CCK and the VIP/non-CCK non-CR populations. The ascending dendrite of the bipolar/bitufted neurons usually finishes in a tuft which starts in L1 in VIP/CR INs and in L2 in the VIP/non-CCK non-CR population, suggesting that the two subtypes may be targeted by different sublaminar L1 inputs ^40^. The axonal arbor also differs in ways that suggest differences in the postsynaptic targets of these populations. Most notably, VIP/CR cells have little axon in the upper part of L2/3 compared to the non-CCK non-CR group which has axon throughout L2/3 (Figure 2 C; Supplementary Figures 2 E, 3 I).

To independently investigate how robustly morphological differences distinguish among the genetically-defined VIP IN populations we performed unsupervised cluster analysis using Ward’s method on 26 well preserved Neurolucida reconstructed VIP neurons recorded in the VIP-Flp; CR-Cre, the VIP-Flp; CCK-Cre and the VIP-Flp; CCK-Cre; CR-Cre; FTLG mice and 6 neurons identified in the VIP-Cre; SNCG-Flp; Ai65 mice. We used 16 morphological parameters: somatic surface area, volume, and location in L2/3; axonal length, surface area and volume; dendritic number, total length, mean length, surface area and volume; and dendritic and axonal x and y vectors; polarity, and sholl analysis. Unsupervised cluster analysis of neuronal morphologies confirmed the presence of four morphological groups that closely matched the molecularly/genetically defined groups (Figure 2E; Supplementary Figures 2-4). The neurons recorded in VIP/SNCG mice clustered with the multipolar neurons recorded in the VIP/CCK mice (clustered together as the same group in later morphological analysis; VIP/CCK H reconstructions from VIP-flp/CCK-cre/FLTG mice n=3; reconstructions from VIP-cre/SNCG-flp mice n=6; total of n=9 VIP/CCK H morphologies). Interestingly, as in the transcriptomic analysis, the multipolar horizontally oriented VIP neurons segregated first from all the VIP IN neurons with bipolar/bitufted morphology, including those in the VIP/CR, VIP/CCK and non-CCK non-CR VIP IN groups. The non-CCK non-CR VIP INs and the bipolar VIP/CCK (VIP CCK V) INs are the most closely related groups in the cluster analysis (Figure 2E). However, we believe they represent two distinct populations. Beyond the difference in expression of CCK in the two groups, the two show statistically significant differences in their axonal and dendritic morphology (Supplementary Figures 2 C&D, 3 G,H,K). The descending dendritic arbor of the bipolar VIP/CCK V INs is proportional to their ascending arbor, while the non-CCK non-CR INs can appear monopolar because their descending dendrite is very short while the ascending dendrite is significantly more prevalent in layer 1 compared to VIP/CCK V (Figures 2 A&B; Supplementary Figures 2 C&D, 3 G,H,K, 4). While the axon of both VIP/CCK V and non-CCK non-CR VIP INs is similar, VIP/non-CCK non-CR Ins have significantly more axon in L5a compared to VIP/CCK V (Supplementary Figure 3 G,H,K). Furthermore, as described below, the two populations differ in their firing patterns and connectivity.

### Distinct intrinsic electrophysiological properties of VIP IN subpopulations

Cortical VIP INs have unique intrinsic electrophysiological properties that distinguish them from other cortical INs. Most notably, VIP INs have a much higher input resistance (Rm) than INs in other IN groups (342 MΩ ^37^) compared to PV INs (69-75 MΩ ^47,48^), SST INs (200 MΩ ^32^; this study), and L2/3 NGFCs (188 MΩ ^49^), a property that makes VIP neurons particularly sensitive to excitatory inputs. Consistent with this, VIP INs require less current injection to reach spiking threshold (rheobase, Rb): 59 pA vs 240-539, 91, and 139 pA for L2/3 PV, SST and NGFCs, respectively. While all VIP IN populations had high input resistance and low rheobase, we found the input resistance was somewhat lower and the rheobase higher in VIP/CR neurons (338±126 MΩ, Mean±SD, n=44; and 63±50 pA, n=26) as compared to the two types of VIP/CCK neurons (VIP/CCK H and VIP/CCK V, respectively, 419 ± 145 MΩ, n=36, and 443 ±167 MΩ, n=22; and 55±31 pA, n=39 and 28±23 pA, n=22) and the VIP/nonCCK nonCR neurons (391±155 MΩ, n=37; and 45±26, n=43; Kruskal-Wallis test followed by a Dunn-Sidak post hoc test: Rm, p=0.0097, VIP/CR vs VIP/CCK H, p=0.0327, VIP/CR vs VIP/CCK V, p=0.0270, VIP/CR vs VIP/nCCK nCR, p=0.4272, VIP/CCK H vs VIP/CCK V, p>0.9999, VIP/CCK H vs VIP/nCCK nCR, p>0.9999, VIP/CCK V vs VIP/nCCK nCR, p>0.9999; Rb, p=0.0006, CCK V vs CCK H, p=0.0010, CCK V vs nCCK nCR, p=0.0367, CCK V vs CR, p=0.0015, CCK H vs nCCK nCR, p>0.9999, CCK H vs CR, p>0.9999, nCCK nCR vs CR, p>0.9999). Interestingly, we also observed that VIP/CCK H neurons had a significantly lower spike threshold than VIP/CCK V neurons (−31 ± 5 mV n=26 versus −38 ± 6 mV n=8, p=0.0052, Mann-Whitney test) and a more depolarized RMP (−61 ± 8 mV n=19 versus −72 ± 7 mV n=6, p=0.0091, Mann-Whitney test).

Studies of VIP INs in cortical slices have also shown that these neurons display variability in their firing patterns in response to step membrane depolarizations, and have firing patterns that are rare among INs in other IN classes, including initial bursting and irregular spiking (IS) ^5,37,50^. We observed similar firing patterns in our intersectional mouse lines and found that this diversity varied among the genetically defined populations (Figure 3). Bursting neurons, characterized by the presence of a depolarizing hump at subthreshold depolarizations near rheobase (likely mediated by T-type Ca2+ channels; ^50^) and a spike burst of 2-3 spikes at >100 Hz riding on the depolarizing hump at rheobase, were much more frequently observed in the VIP/CCK populations. We distinguished three subtypes of bursting (BS) neurons: BS-fast adapting (BS-fAD), resembled the neurons classified as fast adapting (FAD) in ^32,37,50,51^. These neurons were characterized by significant failure in spiking throughout the 1sec depolarizing step during suprathreshold depolarizations. Other BS neurons had slow-adapting repetitive firing during depolarizing steps (BS-sAD). These neurons resembled the cells classified as bNA (bursting non-adapting) in ^32,51^ and the cells classified as HTBNA (high-threshold bursting non-adapting of Pronneke et al., 2015 ^37^). A small percentage of BS neurons resembled the BS-fAD neurons but showed additional spike clusters later during the depolarizing step resulting in a stuttering pattern. All types of bursting neurons were more frequently observed in the VIP/CCK population (∼ 50 % of neurons recorded in VIP/CCK mice had a bursting firing pattern). On the other hand, only 2 (out of 51) neurons recorded in VIP/CR mice and 5 (out of 47) in the non-CCK non-CR VIP population showed a bursting firing pattern.

**Figure 3.**
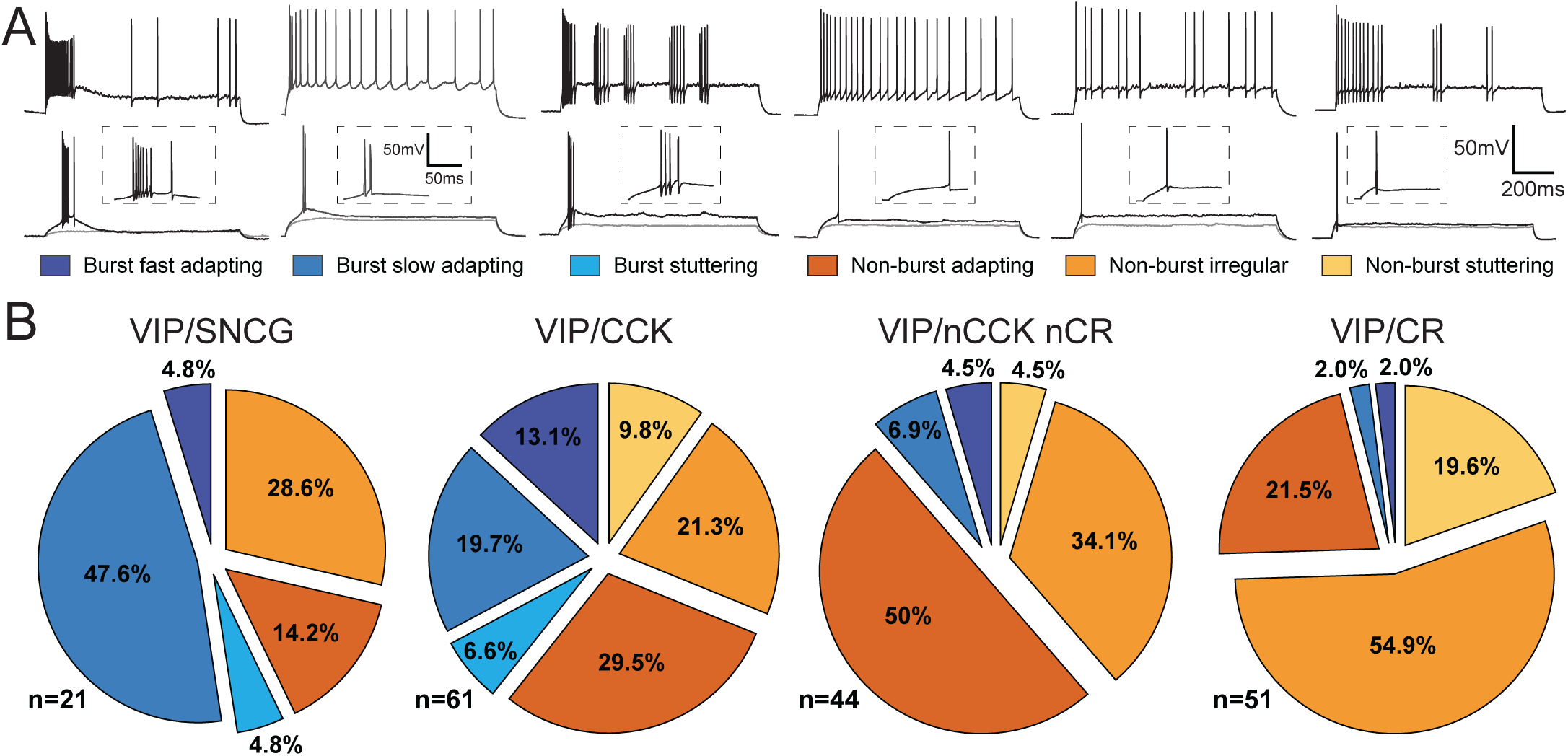
Firing patterns of VIP IN populations. **(A)** Firing properties of VIP INs in response to step depolarizations. Shown are the responses of representative cells to a just subthreshold, threshold, and suprathreshold current injection. Insets show a magnification of the area near the first spike for the threshold response. L2-4 VIP neurons in vS1 showed one of six distinct firing patterns: Burst fast adapting (dark blue) neurons exhibited an initial high-frequency burst of at least three action potentials riding on a calcium hump (inset), followed by rapid adaptation or complete cessation of spiking at rheobase (bottom). During suprathreshold current (top) injections, these neurons largely maintained bursting behavior, though some generated additional spikes. Burst slow adapting (medium blue) neurons fired an initial burst of at least two action potentials at rheobase (bottom; inset), followed by a gradual adaptation in firing rate over the duration of the current injection step at suprathreshold levels (top). Burst stuttering (light blue/cyan) neurons displayed an initial burst of at least three action potentials with fast and low afterhyperpolarizations (AHPs) at rheobase (bottom; inset). During suprathreshold depolarizations, these neurons exhibited intermittent bursts of action potentials with irregular interspike intervals (top). Non-burst adapting neurons (dark orange) generated a single spike at rheobase (bottom; inset) and a train of spikes during suprathreshold current injections, showing a gradual decrease in firing frequency over the course of the depolarizing step (top). Non-burst irregular neurons (medium orange) fired a single spike at rheobase (bottom; inset) and exhibited irregular spiking patterns at suprathreshold depolarization, without a clear adaptation pattern (top). Non-burst stuttering (light orange/yellow) neurons generated a single spike with fast and low AHPs at rheobase (bottom; inset) and displayed intermittent bursts of action potentials with irregular interspike intervals during suprathreshold current injections (top). **(B)** Electrophysiological Subtype Distribution of firing patterns in VIP IN populations. Shown is the proportion of distinct firing patterns for VIP/SNCG (n = 21), VIP/CCK (n = 61), VIP/nonCCK/nonCR (n = 44), and VIP/CR (n = 51) neurons. VIP/SNCG and VIP/CCK neurons predominantly exhibited bursting firing patterns, whereas VIP/nonCCK/nonCR neurons were more commonly associated with non-burst adapting behavior, and CR neurons displayed a higher prevalence of irregular spiking phenotypes.

Among non-bursting neurons we also distinguished three firing patterns. Non BS irregular spiking (non BS-IS), were characterized by the presence of intermittent action potentials elicited with irregular intervals following an initial adapting spike train at near threshold ^51–53^. Non-BS adapting (non-BS AD) resembled the cells classified as CA (continuous adapting of Pronneke et al., 2015 ^37^). Other non-bursting neurons showed stuttering at suprathreshold depolarizations (Non-Burst stuttering). Irregular spiking and stuttering non bursting neurons were more frequently observed among the VIP/CR population, while most non-CCK non-CR neurons had a non-BS AD firing pattern. Among the non-BS neurons in the VIP/CCK population we observed both non-BS AD and non-BS IS neurons at a ∼2:1 ratio. There was no correlation between morphological subtype of VIP/CCK neurons (multipolar or bipolar) and firing pattern, with both subtypes including both bursting and non-bursting firing patterns. Moreover, both types of firing pattern were observed in the VIP/SNCG population, which lacks the VIP/CCK bipolar subtype.

### Differential output connectivity of VIP IN subtypes

The molecular and morphological diversification of VIP INs described above would be physiologically significant if distinct VIP IN subtypes had different patterns of connectivity. There are indications that this is the case. The papers describing the CCK- and CR-expressing VIP neurons in the hippocampus suggested differential connectivity based on anatomical arguments. Most of the axonal arbor of VIP/CR cells in CA1 is located in a sublayer in the stratum oriens/alveus border which is populated by SST INs (OLM cells), but not by PC somas or dendrites, while the axon of VIP/CCK neurons is located in stratum pyramidale, surrounding PC somata in a basket-like manner. EM studies confirmed the differential connectivity predicted by the axonal distribution ^12,13,31^. More recent functional studies confirmed this differential connectivity ^54^. However, in the neocortex, excitatory and inhibitory cells are intermingled. Therefore, such differential connectivity is not expected based on anatomical arguments alone and would require specialized molecular mechanisms.

To study the efferent connectivity of the VIP IN populations, we expressed a channelrhodopsin (CatCh) in VIP/CR and VIP/CCK INs by crossing the intersectional mouse lines with the intersectional CatCh reporter Ai80 (Figure 4A) and used these mice to study the output connectivity of these cells to L2/3 targets in brain slices following light stimulation of the axons of the targeted VIP INs ^5^. We compared the connectivity of these two IN populations to the connectivity of the total VIP IN population using VIP-Flp mice crossed to a Flp-dependent CatCh reporter derived from Ai80 (Ai80F) to compare synaptic responses using the same opsin (Figure 4B). The synaptic responses recorded in these studies likely arise from L2-4 VIP cells. L5-6 VIP cells only have local dendritic and axonal arbors ^37^ and are therefore unlikely to synapse on L2/3 neurons.

**Figure 4.**
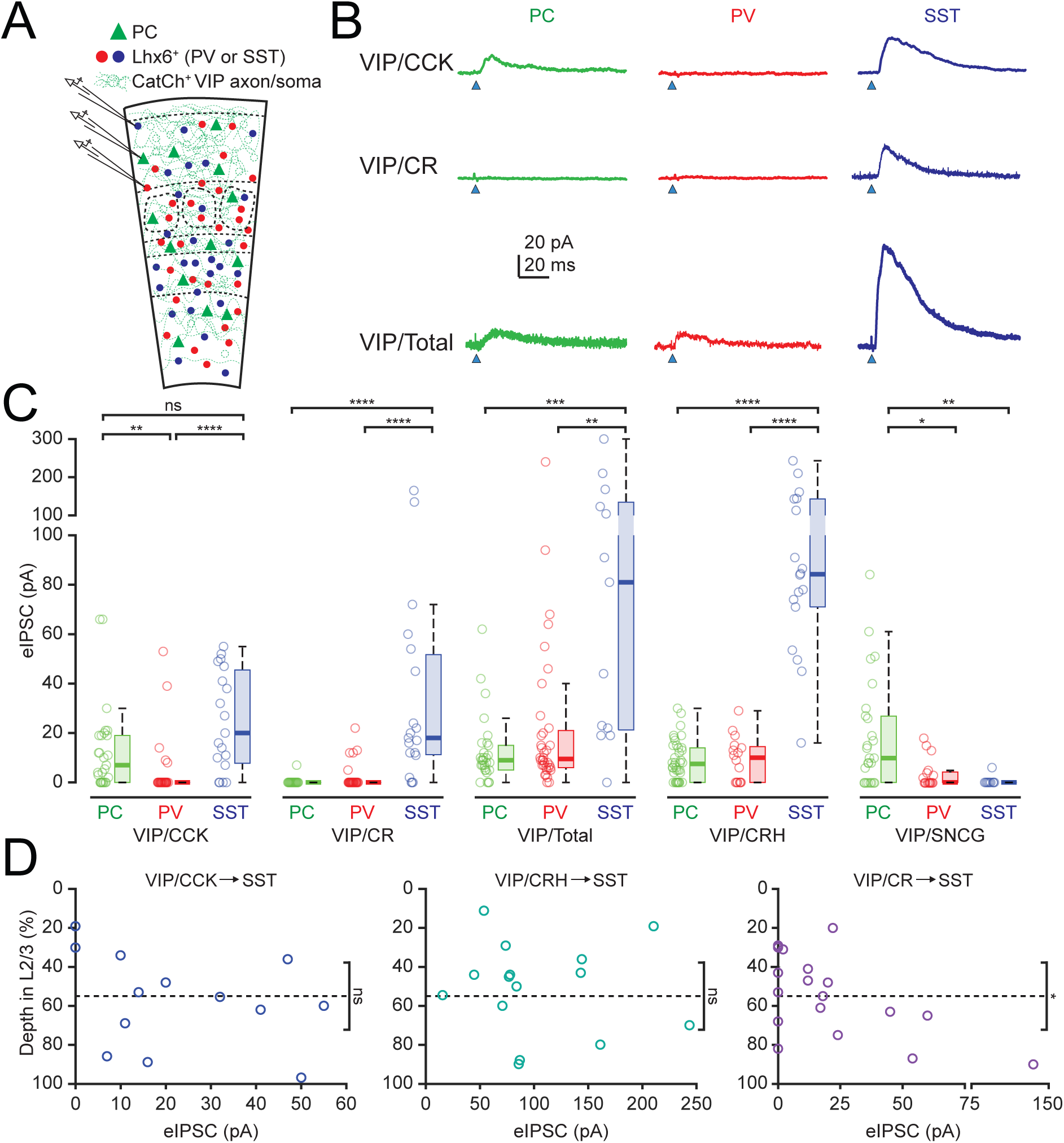
Efferent connectivity of VIP IN populations. **(A)** Schematic of the experiments to study efferent connectivity of VIP IN populations using Ai80 mice. Either PCs or EGFP+ Lhx6+ INs (PV or SST cells) in vS1 were patched to measure output from VIP INs following light stimulation of CatCh-expressing VIP IN axons. **(B)** Representative voltage-clamp (Vhold = −50 mV) recordings of evoked VIP IN output to PCs, PV or SST INs. An unbiased classifier (see Methods) was used to differentiate between PV and SST INs among Lhx6+ neurons. The top, middle, and bottom rows were from representative neurons in the VIP;CCK;Ai80;Lhx6-EGFP, VIP;CR;Ai80;Lhx6-EGFP, and VIP;Ai80-frt;Lhx6-EGFP mouse lines, respectively. **(C)** Quantification of evoked inhibitory postsynaptic currents (eIPSCs) in PCs, PV INs, and SST INs following light stimulation of VIP/CCK, VIP/CR, Total VIP, VIP/CRH and VIP/SNCG IN axons. Box-and-whisker plots the box represents the interquartile range (IQR: 25th to 75th percentile), the horizontal line within the box indicates the median, and whiskers extend to 1.5x the IQR unless outliers are present. Outliers are plotted individually as separate points. (N, respectively, for PC, PV, SST: CCK(27,30,19), CR(22,31,19), Total(30,36,13), CRH(38,16,18), SNCG(17 PC, 10 SST), Kruskal-Wallis test followed by a Dunn-Sidak post hoc test: CCK, p<0.0001, PC vs PV p=0.00371, PC vs SST p=0.32007, PV vs SST p=1.97×10^-5^; CR, p<0.0001, PC vs PV p=0.79165, PC vs SST p=3.42×10^-8^, PV vs SST p=2.74×10^-7^; Total, p=0.0003, PC vs PV p=0.69629, PC vs SST p=0.000203, PV vs SST p=0.002501; CRH, p<0.0001, PC vs PV p=0.99948, PC vs SST p=7.87×10^-9^, PV vs SST p=3.34×10^-6^; SNCG p= 0.0008, PC vs PV p= 0.026569, PC vs SST p= 0.001784, PV vs SST p= 0.57144). **(D)** Quantification of evoked inhibitory postsynaptic currents (eIPSCs) from VIP/CCK, VIP/CRH, and VIP/CR output to L2/3 SST INs by depth in L2/3, with the L1/2 and L3/4 borders being 0% and 100%, respectively. The dashed line indicates the 55% depth mark which is the cutoff for high and low VIP/CR axon density (Figure 2, and supplementary Figure 2). For VIP/CR INs the eIPSC amplitudes in SST INs below this cutoff are significantly larger on average than those above the cutoff (right panel, Two-sample t-test, p=0.0477). In contrast, SST IN eIPSC amplitudes evoked from VIP/CCK or VIP/CRH INs are not significantly different in upper or lower L2/3 (left and middle panels, Two-sample t-test, VIP/CCK p=0.1696 and VIP/CRH p=0.2967).

SST INs are relatively rare in L2/3 (12% of all INs). Therefore, to facilitate the identification of L2/3 PV and SST cells, we incorporated the Lhx6(BAC)-GFP reporter mouse ^55^ which labels all PV and SST cells with GFP, and used their electrophysiological features to distinguish between the two cell types using a classifier with 94.2% accuracy (Supplementary Figure 5).

**Figure 5.**
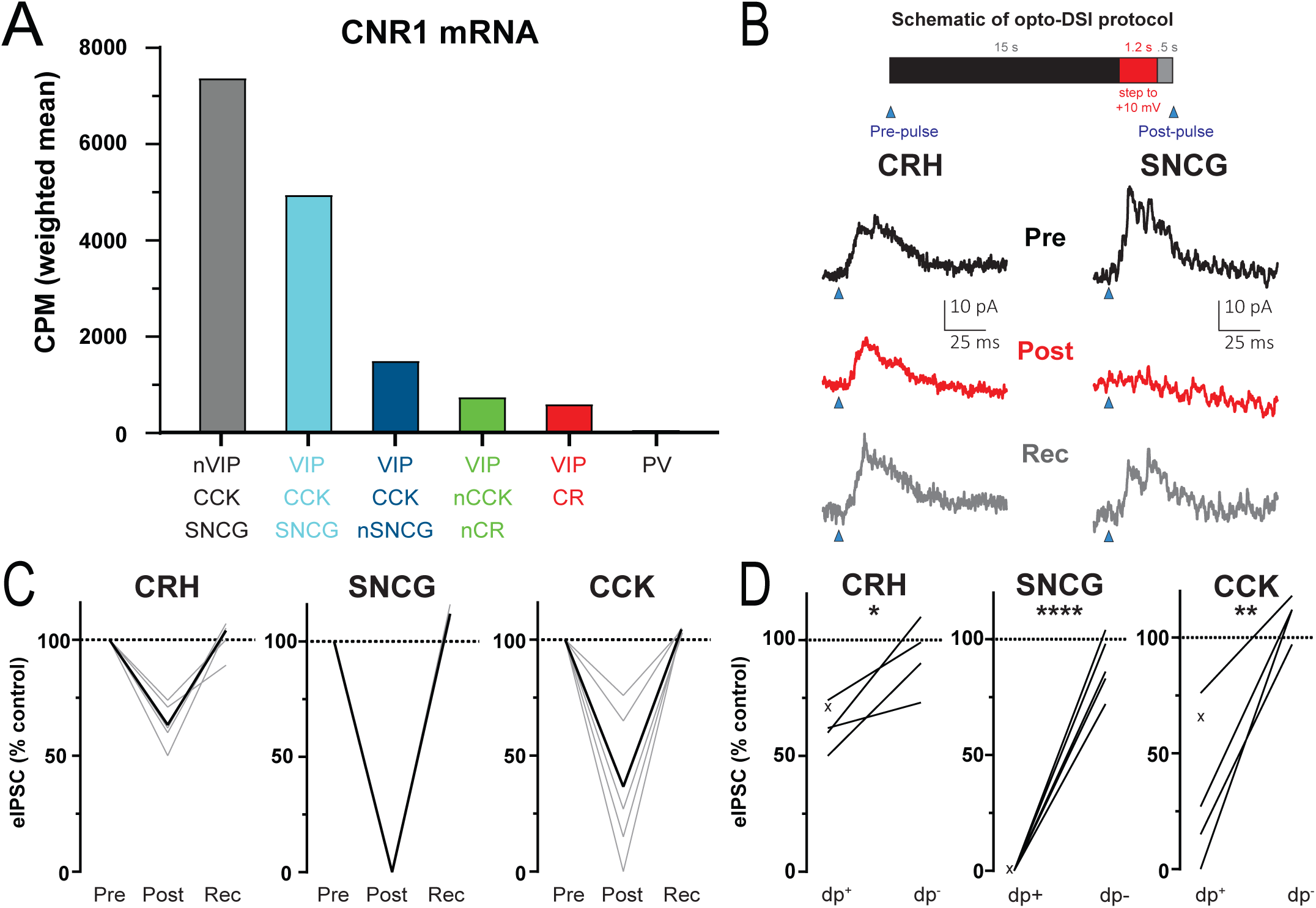
DSI of synaptic responses evoked by VIP INs in L2/3 PCs. **(A)** Stacked bar plots of normalized mRNA levels (CPM weighted mean, the mean of all CPM values from all transcriptomic bins for the indicated IN subtype weighted by the number of cells in each transcriptomic bin) from the Allen Institute library for cannabinoid receptor 1 (CNR1) (Allen Institute scRNAseq data;^35^) in different VIP IN populations and PV neurons. **(B)** Protocol used to study depolarization-induced suppression of inhibition (DSI) (top). DSI was assessed using optogenetics in L2/3 pyramidal cells (PCs) as follows: PCs were recorded in voltage-clamp mode (V_hold_ = −50 mV) in cortical slices from VIP-Flpo/CRH;Ai80, VIP-Cre/SNCG-Flpo;Ai80 or VIP-Flpo/CCK-Cre;Ai80 mice, in which CatCh is expressed in the intersected VIP IN subpopulation. Control IPSCs were elicited by light pulses of 2 ms duration. Then, after a 15 s interval, the PC was depolarized (dp) to +10 mV (from −70 mV) for 1.2 s, and after a 0.5 s recovery, another IPSC was elicited by light stimulation. Averaged traces (n=3) from single cells of pre-DSI (black; top), post-DSI (red; middle), and recovery (bottom; grey) from VIP/CRH (left) and VIP/SNCG (right) Ai80 mice. **(C)** DSI of all cells tested with responses normalized to the pre-depolarization step. Grey, individual cells; black, average of all cells (CRH, n=5; SNCG, n=6; CCK, n=5; Wilcoxon matched-pairs signed rank test, one-tailed, post vs pre, CRH, p=0.0312, SNCG, p=.0156, CCK, p=.0312). The magnitude of DSI observed in SNCG was significantly greater than that observed in CRH (Kruskal-Wallis test, p=0.002 followed by a Dunn-Sidak post-hoc test, SNCG vs. CRH, p=0.0071, SNCG vs. CCK, p=0.0894, CRH vs. CCK, p>0.9999), **(D)** Summary of all cells, comparing for each cell, the effect of withholding the depolarization step (dp^+^ vs dp^-^, Wilcoxon matched-pairs signed rank test, one-tailed, SNCG, p=0.0312; Paired t-test, one-tailed, CRH, p=0.0173, CCK, p=0.0057). Cells in which no dpprotocol was run are denoted by an ‘x’.

The responses observed when the total VIP IN population was stimulated in VIP-Flp mice were similar to those reported previously with VIP-Cre mice in vS1 ^5^ as well as in V1, auditory and medial prefrontal cortices ^8,9^. Synaptic responses were largest on SST INs, and they were small but significant on PCs and PV INs (Figure 4C). In contrast, light activation of VIP/CR IN axons in VIP-Flp; CR-Cre; Ai80 mice essentially produced no synaptic responses on PCs or PV cells, but produced substantial responses on SST cells, even on SST cells recorded simultaneously with a nearby non-responsive PC. On the other hand, light activation of VIP/CCK cell axons produced responses in PCs, comparable to those on SST cells). As with VIP/CR INs, there were negligible responses on PV cells (Figure 4C). The magnitude of the synaptic responses elicited by VIP/CCK IN stimulation on PCs was similar to that elicited when the total VIP IN population was stimulated (Figure 4C), suggesting that *most if not all* the connectivity of VIP INs with PCs is derived from VIP/CCK cells, and that the non-CCK non-CR VIP INs, like VIP/CR neurons, don’t synapse significantly on PCs. On the other hand, the lack of connectivity of both the VIP/CCK and the VIP/CR INs to PV cells suggests that this connectivity is produced by the non-CCK non-CR VIP population.

To obtain more direct information on the connectivity of the non-CCK non-CR VIP INs, we searched the single cell transcriptomic data from the Allen Institute’s cortical scRNAseq collection ^35^ to identify genes that could be used to target this IN population. We found two genes that are expressed in non-CCK non-CR VIP INs and are not expressed in VIP/CR neurons: Cxcl14 and CRH (Supplemental Figure 1). However, both are also expressed in some of the VIP/CCK neurons. Of the two genes, we reasoned that CRH would be more useful, because it was not expressed in the VIP/CCK INs of the SNCG cluster (Supplementary Figure 1). Thus intersectional VIP/CRH mice (VIP-Flp; CRH-Cre; Ai80) could be used to independently infer the connectivity of the non-CCK non-CR VIP population and to distinguish between the two VIP/CCK IN populations. Light stimulation of slices from VIP-Flp; CRH-Cre; Ai80 mice produced large synaptic responses on L2/3 SST neurons (Figure 4C). We also observed synaptic responses in slices from these mice on PV neurons, consistent with the prediction that connectivity of VIP INs to PV INs arises from the non-CCK non-CR VIP population. We also observed responses on PCs, similar to those observed on VIP/CCK INs. We hypothesize that these synaptic responses arise from stimulation of the axons from the VIP/CCK neurons labeled in the VIP/CRH intersectional mouse line, based on our prediction that the non-CCK non-CR VIP INs have negligible connectivity to PCs (see above). Additional evidence in support of the view that the connectivity with PCs observed in the VIP/CRH line arises from the VIP/CCK neurons present in this line is presented below.

The VIP/SNCG neurons, which are excluded in the VIP-Flp; CRH-Cre intersectional cross are also expected to synapse on PCs based on the conclusion that they correspond to the small CCK basket cells (see above). This was confirmed by light stimulation of slices from VIP-Cre; SNCG-Flp; Ai80 mice (Figure 4C). These observations suggest that the two VIP/CCK IN populations, the vertically oriented VIP/CCK V neurons and the horizontally oriented VIP/CCK H neurons connect to L2/3 PCs. Interestingly, VIP/SNCG neurons had negligible connectivity to SST INs (Figure 4C), indicating that the connectivity to these INs in VIP/CCK mice arises entirely from the vertically oriented VIP/CCK INs.

Both the non-CCK non-CR VIP INs and the VIP/CR INs connected preferentially to SST INs, however, the synaptic responses on SST cells from non-CCK non-CR VIP cells were considerably larger than those from VIP/CR neurons (Figure 4C). Morphological analysis showed that while non-CCK non-CR VIP INs have a high axonal density throughout L2/3, the upper part (upper 55%) of these layers is essentially devoid of axon from VIP/CR INs (Figure 2; Supplementary Figure 2-3). We therefore compared the synaptic responses of L2/3 SST INs elicited by light stimulation of VIP IN axons as a function of their location in L2/3 (Figure 4D). Consistent with the axonal distribution of VIP/CR cells, the response of SST INs located in the upper part of L2/3 to the stimulation of VIP/CR axons was significantly smaller than the response of SST INs located in the deep part of L2/3. In contrast, there was no difference in the response of superficial or deep L2/3 SST INs to stimulation of VIP/CRH and VIP/CCK axons.

### Differential synaptic properties of the two VIP/CCK IN populations

We examined the Allen transcriptomic data ^35^ for molecular differences between the two types of VIP/CCK INs and found that they differ in terms of expression of the CB1 endocannabinoid receptor (Cnr1), which is highly enriched in the multipolar/horizontal VIP/SNCG neurons as compared to the bipolar/vertical VIP/CCK V cells (Figure 5A). We investigated whether this resulted in differences in synaptic connections to PCs. Endocannabinoids are known to mediate a phenomenon known as depolarization induced suppression of inhibition (DSI; ^56,57^, which has been observed on the synaptic connections of CCK BCs ^58,59^. Depolarization of a connected pyramidal cell leads to the release of endocannabinoids from the pyramidal cell that then bind to CB1 receptors located on the pre-synaptic terminals of the CCK BCs, leading to suppression of GABA release ^60–62^. Consistent with the difference in their levels of expression of CB1 receptors, we observed a much stronger DSI on the synaptic responses elicited by light stimulation of the VIP/CCK H INs in the VIP/SNCG mouse as compared to light stimulation of the VIP/CCK V INs and the non-CCK non-CR VIP neurons in the VIP/CRH mouse (Figure 5B-D). The DSI of the responses elicited on VIP/CCK mice, in which all VIP/CCK INs are stimulated, was intermediary, between to those observed on VIP/SNCG and VIP/CRH mice (Figure 5C&D). This result supports the conclusion that both types of VIP/CCK INs target PCs.

### Differential input connectivity of VIP IN subtypes

The studies of efferent connectivity show that distinct VIP IN subtypes differ in their postsynaptic targets. This would be highly significant if these populations were driven by distinct excitatory sources, implying that different behavioral contingencies recruit distinct VIP IN subtypes and thereby inhibit or disinhibit PCs, or inhibit different subtypes of SST INs and thereby disinhibit distinct types of PCs. We therefore next investigated the input connectivity of the VIP IN populations (Figure 6). Of the three major long-range inputs to vS1: vibrissa motor cortex (vM1), secondary somatosensory cortex (S2) and the somatosensory higher-order thalamic nucleus POm, the inputs from vM1, which are thought to mediate sensorimotor integration in the whisker system, have been shown to preferentially target VIP INs in L2/3 ^5,63^. To explore the connectivity of vM1 inputs to distinct VIP IN subtypes in vS1, we virally expressed ChR2 in vM1 excitatory neurons ^5^ and recorded postsynaptic responses to light stimulation in identified VIP IN populations of L2-4 of vS1 (Figure 6 A-F). All VIP IN subtypes responded to light stimulation of vM1 afferents, but the response was significantly larger on the non-CCK non-CR VIP population, as compared to the VIP/CCK V and VIP/CR populations, and often led to action potential firing when recorded under current clamp (Figure 6 E&F). Synaptic responses were also smaller on VIP/CCK H neurons than in the non-CCK non-CR VIP population, but this difference was not statistically significant.

**Figure 6.**
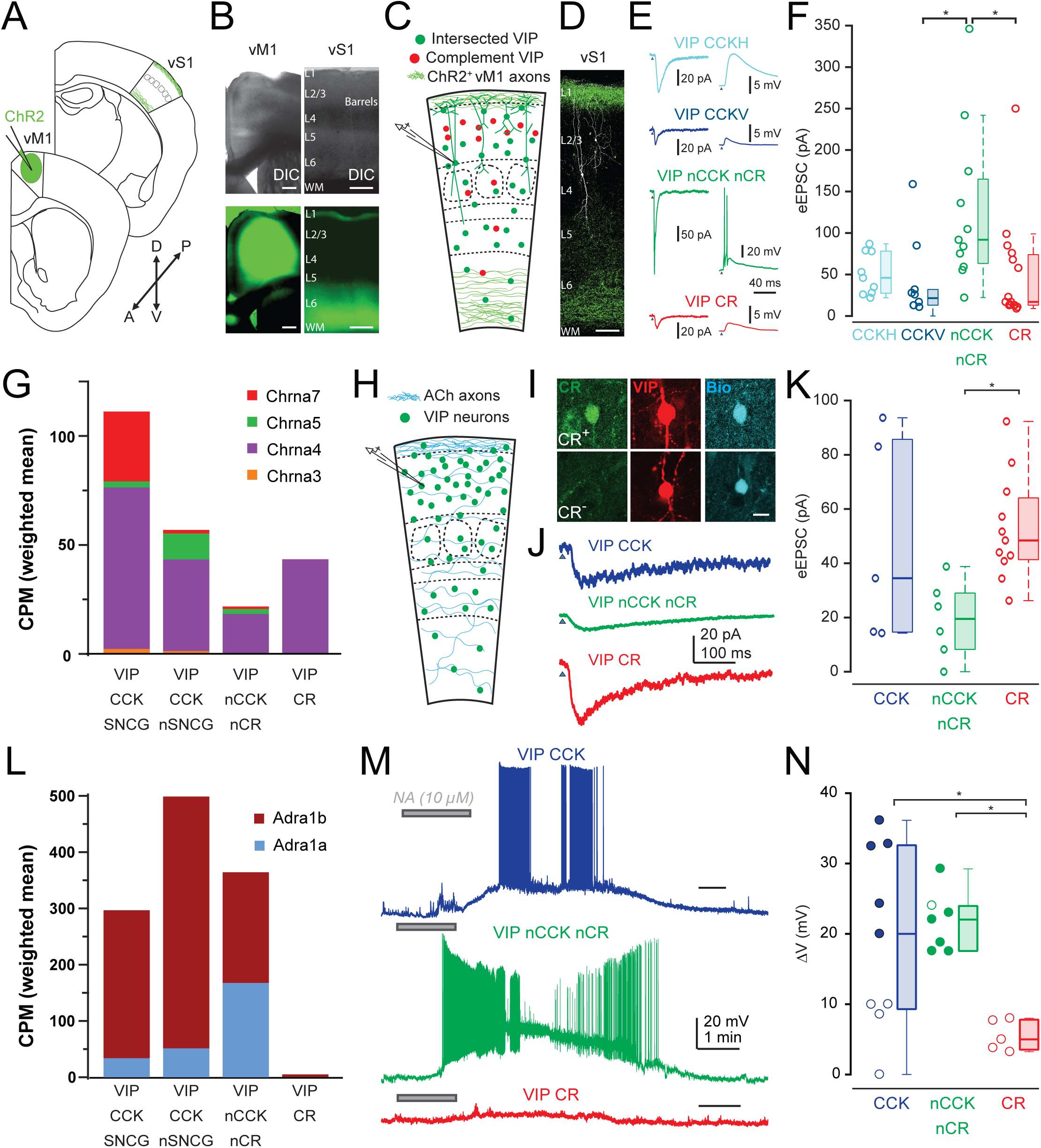
Differential afferent connectivity of VIP IN populations. **(A)** Schematic representation of the experimental procedure to study vM1 inputs onto L2-4 VIP INs in vS1, showing the injection of AAV-ChR2-EYFP into vM1, leading to the expression of ChR2 in vM1 axons projecting to Layer 1 and Layer 6 of vS1. **(B)** Differential interference contrast (DIC) (top panels) and epifluorescence images (bottom panels) of the vM1 injection site (left) and the projection/recording site in vS1 (right), illustrating the ChR2-EYFP injection site and the vM1 axons in vS1. Scale bar = 200 μm. **(C)** Schematic of experimental procedure. VIP/SNCG, VIP/CCK, VIP/nonCCK nonCR, or VIP/CR neurons were patched in the appropriate mouse lines and ChR2 was expressed in vM1 in order to optically stimulate vM1 projections in vS1. **(D)** Confocal image showing two patched VIP neurons, with a VIP/CR neuron (left, white) and a VIP/CCK neuron (right, white), both receiving vM1 input from L1-projecting axons (green top). Scale bar = 200 μm. **(E)** Representative voltage-clamp (left, V-clamp, V_hold_ = −90 mV) and current-clamp (right, I-clamp, V_hold_ = −70 mV) recordings from the samecells of vM1 evoked synaptic input to VIP/CCKH (cyan), VIP/CCKV (blue), VIP/nCCK nCR (green), and VIP/CR (red) neurons. **(F)** Quantification of vM1 evoked excitatory postsynaptic currents (eEPSCs) in different VIP IN populations (CCK H N=15, CCK V N=10, nonCCK nonCR N=11, CR N=15). Box-and-whisker plots. The box represents the interquartile range (IQR: 25th to 75th percentile), the horizontal line within the box indicates the median, and whiskers extend to 1.5x the IQR unless outliers are present. Data points which lie beyond the range of the whiskers are considered outliers. The vM1 response evoked in nCCK nCR VIP INs was significantly stronger than VIP/CCK V and VIP/CR (Kruskal-Wallis p = 0.042, Dunn-Sidak nCCK nCR vs CCK V p = 0.0166 and nCCK nCR vs CR p = 0.0191). The VIP/CCK H group received the second strongest input and was not significantly different from the rest of the groups (CCK H vs CCK V p = 0.2037, CCK H vs nCCK nCR p = 0.8228, CCK H vs CR p = 0.2682) and the least responsive groups were VIP/CCK V and VIP/CR (CCK V vs CR p = 0.9997). **(G)** Stacked bar plots of normalized mRNA levels (CPM weighted mean, the mean of all CPM values from all transcriptomic bins for the indicated IN subtype weighted by the number of cells in each transcriptomic bin; Allen Institute scRNAseq data;^35^) for nicotinic cholinergic receptor α subunits (Chrna3, Chrna4, Chrna5, Chrna7) across four VIP IN populations (VIP/CCK SNCG, VIP/CCK nSNCG, VIP/nonCCK nonCR, and VIP/CR) showing that although all VIP populations express α nicotinic receptor subunits the VIP/nonCCK non CR neurons expressed the lowest levels. **(H)** Schematic of the cholinergic input experiment, where L2-4 VIP INs were patched in a Cholinergic/VIP mouse model (ChAT-ires-Flpo; VIP-Cre; Ai80F + Cre-dependent tdTomato virus to label all VIP INs) in Layer 2/3, and measuring cholinergic light-evoked currents in the presence of synaptic blockers (Gabazine and CNQX 10 µM; D-AP5 25 µM). Soma location, morphology, electrophysiological properties, and CR immunohistochemistry were used for VIP IN classification. **(I)** Post hoc immunohistochemistry in brain slices showing biocytinfilled patched neurons. The images show CR+ (top) and CR-(bottom) neurons, co-labeled with VIP (red) and Biocytin (blue). Scale bar = 10 μm. **(J)** Representative voltage-clamp (V-clamp) recordings of cholinergic light-evoked currents in VIP/CCK (blue), VIP/nonCCK nonCR (green), and VIP/CR (red) neurons. **(K)** Quantification of cholinergic input (eEP-SCs) to VIP neurons (CCK, N=5, nonCCK nonCR N=6, CR N=11, Kruskal-Wallis test p=0.0127, Dunn-Sidak CCK vs nonCCK nonCR p=0.3129, CCK vs CR p>0.9999, nonCCK nonCR vs CR p=0.0147). **(L)** Stacked bar plots of normalized mRNA levels (CPM weighted mean as in **G**; Allen Institute scRNAseq data;^35^) for α1-adrenergic receptor subunits (Adra1a, Adra1b) across the four VIP interneuron subpopulations (VIP/CCK SNCG, VIP/CCK nSNCG, VIP/nonCCK nonCR, VIP/ CR). The height of each colored segment represents the relative contribution of each subunit to the total expression. **(M)** Representative traces of noradrenaline (NA, 10 μM; Tocris 5169) application in the presence of glutamatergic blockers (D-AP5 25 μM; Abcam and CNQX 10 μM; Abcam), showing differential effects on a VIP/CCK, VIP/nonCCK nonCR, and a VIP/CR IN. The VIP/CR neuron exhibited a modest membrane potential depolarization, whereas the VIP/CCK and VIP/nonCCK nonCR INs both showed a strong depolarization with spiking. **(N)** Quantification of membrane potential change (ΔV, if cell began spiking than spiking threshold was used as the final Vm reached; cells that spiked are marked as filled in circles, otherwise they are empty) after NA application demonstrating a significantly greater NA-induced depolarization in VIP/CCK and VIP/nonCCK nonCR neurons, with many reaching spiking threshold, compared to VIP/CR neurons (CCK, N=9, nCCK nCR N=7, CR N=5, Kruskal-Wallis test p=0.0091, Dunn-Sidak CCK vs nCCK nCR p>0.9999, CCK vs CR p=0.0336, nCCK nCR vs CR p=0.0225).

VIP INs are also strongly stimulated by cholinergic stimulation, likely mediated primarily by α4β2 nicotinic receptors ^64^. Stimulation of ChAT cells or arousal produces much larger non-glutamatergic excitation of VIP INs than any other cells in L2/3 (Figures 1E, 3C in ^65^), but the responses of individual cells are highly variable. This variability may arise from differential targeting of VIP IN subtypes by cholinergic inputs. Transcriptomic analysis shows that all VIP IN populations express several nicotinic receptor alpha subunits, with α4 being the dominant subtype (Figure 6 G). The levels of α4 transcripts were lower in the non-CCK non-CR VIP INs as compared to VIP/CCK and VIP/CR subtypes (Figure 6 G). Consistent with this pattern, the responses elicited by light stimulation of ChR2-expressing cholinergic axons were lower in the non-CCK non-CR VIP INs than those recorded on L2/3 VIP/CCK or VIP/CR neurons (Figure 6 H-K).

Transcriptomic analysis ^35^ also shows differential expression of α1a and α1b adrenergic receptors of VIP IN subtypes (Figure 6 L) and VIP INs have been shown to be synaptically coupled to NA neurons and respond to NA input ^66–68^. Transcriptomic subtypes that correspond to VIP/CCK INs strongly express α1b receptors, while those that correspond to the non-CCK non-CR VIP population express both α1a and α1b. Adrenergic receptors were only weakly expressed in VIP/CR INs. We compared the effect of bath application of the adrenergic agonist noradrenaline (10 µM) on VIP/CR, VIP/CCK and VIP/non-CCK non-CR neurons. Consistent with predictions based on transcriptomic data, responses were significantly smaller on VIP/CR cells compared to those on the VIP/CCK and VIP/non-CCK non-CR populations (Figure 6 M&N).

### VIP IN-mediated inhibition and disinhibition in neocortex

The connectivity data suggest that while the activity of VIP/CR INs is likely to produce disinhibition of PCs, activity in VIP/CCK cells might inhibit PCs, since their relatively weaker connection to INs might not be sufficient to produce disinhibition. In fact, most INs connect to other INs ^3,9^. To produce disinhibition of PCs, the reduced inhibition from SST cells has to be able to overcome the inhibition the PCs receive directly from the VIP/CCK cells or other INs active in the same context.

We used silicon probes to record the effects of optogenetic stimulation of VIP/CR or VIP/CCK INs on the activity of PCs in freely moving mice (Figure 7A). We used the intersectional VIP/CR and VIP/CCK mice crossed to the intersectional CatCh reporter mouse Ai80. Light stimulation of VIP/CR INs increased the activity of a fraction of the recorded PCs, and inhibited a population of INs, presumably SST cells (Figure 7 B-D). On the other hand, light stimulation of VIP/CCK INs inhibited a fraction of the recorded PCs (as well as some INs) (Figure 7 E-G). The observed effect is likely an underestimate of the inhibition of PCs by VIP/CCK INs, since silicon probe recordings are biased against superficial layers ^49,69^, where the axon of VIP/CCK cells is concentrated. These results support the hypothesis that VIP/CR INs are disinhibitory while VIP/CCK neurons may predominantly produce inhibition as predicted by the connectivity studies in slices (Figure 7 H).

**Figure 7.**
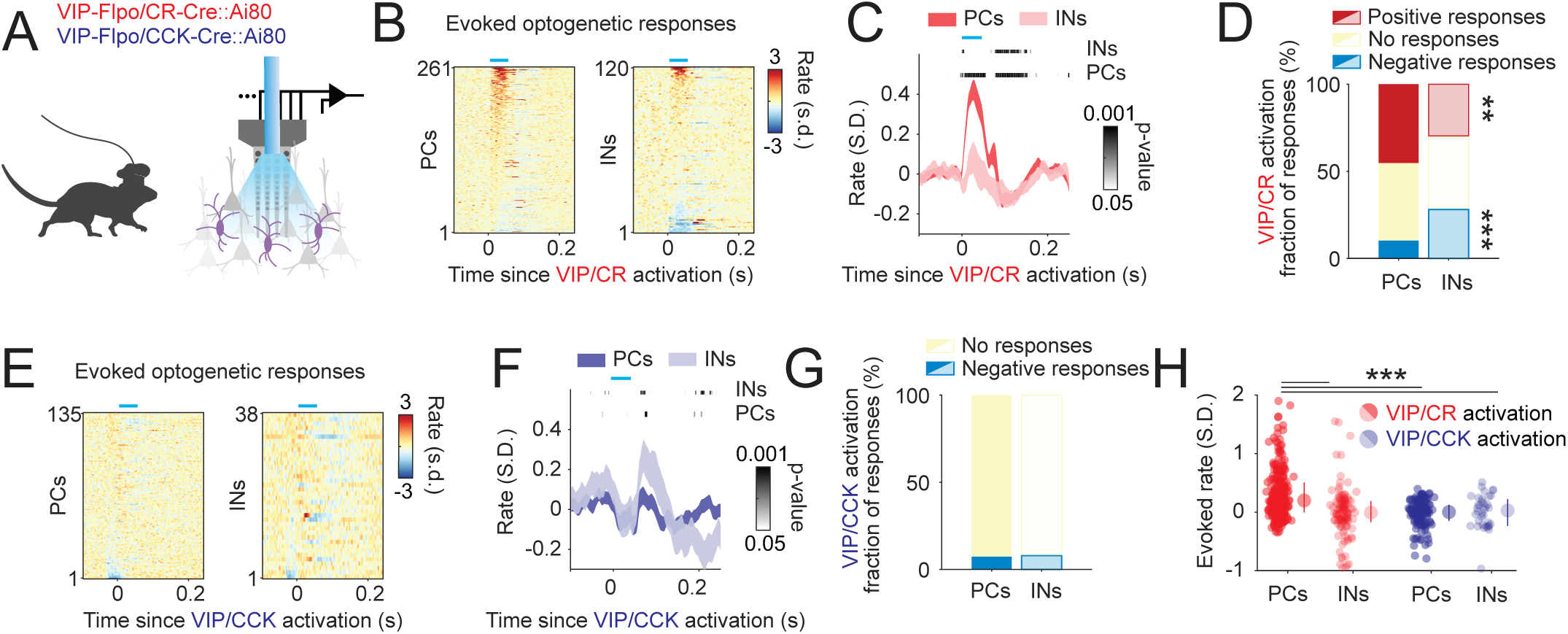
Network effects of the activation of VIP/CR and VIP/CCK INs. **(A)** Optogenetic activation of VIP/CR and VIP/CCK neurons in freely behaving mice. Combined optical fiber-electrode probes were implanted in VIP-Flpo; CR-Cre; Ai80 and VIP-Flpo; CCK-Cre; Ai80 mice to enable simultaneous stimulation and recording. **(B)** Peristimulus time histograms (PSTHs) of pyramidal cells and interneurons in the posterior parietal cortex (PTLp) following VIP/CR IN activation, sorted by light stimulus (n = 4 mice). **(C)** Average firing rate (± s.d.) peri-stimulus time histograms (PSTHs) for pyramidal cells and interneurons. The significance of the average response in each spatial bin was assessed using the Wilcoxon rank test and is shown for both groups. **(D)** Fraction of positive, negative, and non-significant responses (bootstrap, 1000 repetitions) following VIP/CR IN activation across groups. A higher fraction of pyramidal cells was disinhibited by VIP/CR optogenetic activation (45.2% vs. 30.0%, *p* = 0.005, Chi-square test). Conversely, a greater fraction of interneurons was suppressed by the same manipulation (10.3% vs. 28.3%, *p* < 10^-5^). **(E-F)** Same as in B-C for VIP/CCK IN activation (n = 2 mice). **(G)** A comparable fraction of pyramidal cells and interneurons was suppressed by VIP/CCK IN activation (7.4% and 7.8%, p = 0.95). **(H)** Pyramidal cells were disinhibited by optogenetic activation of VIP/CR INs (two-way ANOVA: effect of optogenetically activated cell group, *F_1,549_* = 16.5, *p* < 10^-4^; effect of response group, *F_1,549_*= 8.2, p = 0.004; interaction between activated and response groups, *F_1,549_* = 11.8, p < 10^-3^).

## Discussion

The development and introduction of genetic methods in rodents to label and manipulate the major IN populations in cortex (PV, SST, VIP and ID2 or Lamp5) have been transformative, and have resulted in major advances in our understanding of cortical circuits ^70,71^. It is now evident that these IN groups are heterogeneous and are comprised of discrete subtypes that often have important differences in connectivity and function. For example, PV INs encompass both basket cells and chandelier cells, populations that share similarities in molecular gene expression and intrinsic electrophysiological properties but that have different output connectivity: basket cell terminals target the soma and proximal dendrite of PCs whereas axo-axonic cells target the PC axon initial segment. Recent work has underscored the importance of this next level of organization, as illustrated by the monumental single-cell transcriptomic analyses of cortical neuron types by the Allen Institute ^33–35,39^, the analysis of the diversity of PV basket cells ^72–74^, of chandelier cells ^75^, and of SST INs ^76–81^. Linking the transcriptomic diversity of IN subtypes to their functional properties, circuit function and *in vivo* activity has begun to be achieved via remarkable post-hoc efforts ^33,81–85^, but this remains a dauntingly complex endeavor. Considering that no single non-overlapping set of genes have been identified for the vast majority of IN subtypes, progress in understanding the functional specialization of IN subpopulations has benefitted from the development of new genetic approaches that give experimental access to this diversity in a practical and reasonable way.

In this study we applied intersectional genetic methods to reveal clearly distinct functional populations of VIP INs (Figure 1). We found that VIP INs can be subdivided into four main groups that exhibit distinct laminar organization, stereotypical patterns of axonal and dendritic arborization and differences in input and output connectivity (Figures 2&4). The two types of VIP/CCK INs, which account for about 30% of L2-4 VIP neurons both provide inhibition onto PCs (Figure 4 C), but had interesting differences in their synaptic properties (Figure 5) as well as their transcriptomic profiles (e.g., Ca^2+^ related genes such as Cadps2; Supplementary Figure 1). They are predicted to mediate inhibition on PCs while the VIP/CR cells and the non-CCK non-CR VIP INs predominantly target SST INs and are predicted to be disinhibitory (Figure 4 C). *In vivo* experiments exploring the effect on the cortical network of light stimulation of ChR2 expressing VIP/CCK or VIP/CR neuron subtypes in freely moving mice provided support to these predictions (Figure 7).

Interestingly, although both the VIP/CR and the non-CCK non-CR VIP INs predominantly target SST INs, we found that they differed in the populations of L2/3 SST INs that receive strong innervation from each subtype. VIP/CR INs mainly target SST INs in deep L2/3, while the non-CCK non-CR VIP INs equally targeted superficial and deep L2/3 SST INs (Figure 4). Distinct SST IN subtypes, with different localization in the cortical column have been shown to inhibit different types of PCs and/or different compartments of the same PC ^80^. This predicts that distinct VIP IN subtypes may disinhibit distinct subsets of PCs. Moreover, we found that non-CCK non-CR VIP neurons but not VIP/CR cells also target PV INs, albeit much less than SST INs, suggesting that inputs targeting these VIP INs produce disinhibition of both the soma and the distal dendrite of PCs, and thus could facilitate both feedforward and feedback inputs (Figure 4).

Examination of the inputs to VIP INs in vS1 by long-range projections also indicated that there is differential targeting of VIP subtypes. Glutamatergic inputs from vM1 preferentially targeted non-CCK non-CR VIP INs, whereas these VIP neurons received the weakest input from cholinergic axons from the basal forebrain (Figure 6 A-K). Lastly, we found strong noradrenergic responses on both VIP/CCK and non-CCK non-CR VIP INs but not on VIP/CR cells (Figure 6 L-M). Given their differential efferent connectivity, this indicates that the functional effect of any stimulus that activates VIP INs depends on which VIP IN subpopulations are targeted ^86^. Together, the results suggest that VIP IN subtypes form highly specific circuits, as has been proposed for SST IN subtypes ^80^. While the molecular basis for the differential input and output connectivity of VIP IN subtypes is not yet clear, there are some notable cell surface molecules that show selective expression in specific populations (e.g., the protocadherin Pcdh11x in non-CCK non-CR VIP INs; Supplementary Figure 1). Considering recent evidence for distinct VIP synaptic dynamics on different SST IN subtypes^87^, it will be interesting to elucidate how the molecular heterogeneity of VIP and SST subpopulations results in specialized disinhibitory subcircuits.

L2-4 VIP INs project a significant amount of axon to deep cortical layers (Figure 2 and Supplementary Figures 2 C-E, 3 F-I, K), where a diverse population of SST INs reside ^3,76–80^. Furthermore, paired recordings have shown that L2/3 VIP cells strongly connect to deep layer SST cells ^9,88,89^. It will be of great interest to investigate whether L2-4 VIP/CR and non-CCK non-CR VIP INs differentially target distinct L5-6 SST subpopulations. This requires the development of methods to identify specific SST subtypes that are compatible with the genetics used to select distinct VIP IN subtypes for stimulation ^90^. The recent discovery of enhancers labeling specific SST IN subpopulations is promising in this regard ^91^. These studies will also require the ability to express optogenetic actuators in VIP INs in superficial cortical layers without contamination from VIP INs in deep layers.

The VIP-SST disinhibitory circuit motif has emerged as a key circuit mechanism for top-down modulation and context-dependent cortical processing (see recent reviews by ^3,5,15,92–99^. The existence of VIP IN subtypes with different output connectivity may explain some results that appear to contradict the view that increased activity of VIP neurons produces disinhibition of excitatory neurons ^100^. For instance, while in V1, locomotion increases sensory responses ^94^, in auditory cortex (A1) it does the opposite, although in both cases VIP INs are recruited. The VIP disinhibitory circuit has been implicated as the cause of increased responses in V1 ^94^. A recent study, while confirming that movement suppresses responses in auditory cortex, shows that in contrast, optogenetic activation of VIP INs in this cortical area also increases stimulus-evoked spike rates ^101^. One possibility is that while activating VIP INs optogenetically excites all VIP subtypes, in A1, movement might preferentially recruit inhibitory VIP/CCK cells. In another example Batista-Brito (2017) ^21^ showed that deletion of the schizophrenia-related gene ErbB4 specifically from VIP INs impaired VIP IN function and produced dramatic effects on visual responses and severe deficits in cortical activity, visual responses, sensory perceptual function and other behavioral changes characteristic of models of schizophrenia. However, deletion of ErbB4 from VIP-INs also led to a marked increase in the activity of cortical excitatory neurons. One possibility is that ErbB4 deletion in VIP cells primarily affected VIP/CCK INs, and that it is the impairment of VIP IN-mediated inhibition that is responsible for the effects.

There is increasing evidence for heterogeneity of VIP IN activity *in vivo*. A recent study investigating stimuli encoding in visual cortex reported the presence of distinct subpopulations of VIP INs (∼5) encoding specific combinations of novel stimuli, omissions of familiar stimuli or behavioral outcomes in an image change detection task ^102^. It is possible that this observed heterogeneity is related to the diversity of VIP IN populations revealed by this study or via transcriptomic studies. Differences in intrinsic electrophysiology, connectivity, and response to neuromodulators, as observed in this study, could all affect the respective *in vivo* activities of VIP subpopulations ^103^. In sum, it is increasingly clear that the heterogeneity of VIP INs observed in molecular profiling and genetic studies is functionally important, and provides additional flexibility and modularity to disinhibitory circuitry in the brain.

## Resource Availability

Requests for further information and resources should be directed to and will be fulfilled by the lead contact, Bernardo Rudy.

All unique reagents, code, and data generated in this study are available from the lead contact upon request.

## Acknowledgments

We thank Chiung-Yin Chung for her superlative lab and mouse colony management; Maximiliano Jose Nigro, Brian Clark, Sebnem Tuncdemir, and Robin Tremblay for their data to train the PV/SST classifier; Benjamin Schuman and Bryce Grier, and all members of the Rudy Lab for their helpful discussions and guidance; Gord Fishell and Chris Harvey for providing viral constructs S9E10 and S44 respectively to label SST neurons. This work was supported by NIH grants R01NS133751 (to BR), R21MH137452 (to BR), P01NS074972 (to BR), and R01MH062349 (to XJW).

## Author Contributions

Conceptualization and methodology, S.D., H.Z., M.V., E.G., A.P., J.L.H., R.M., B.R.; investigation, S.D., H.Z., I.K., M.V., P.A., E.G., A.P., J.L.H., E.M., M.O., R.M.; analysis, S.D., H.Z., M.V., P.A., J.M., J.L.H., R.M.; writing – original draft, B.R.; writing – review and editing, S.D., H.Z., R.M., B.R.; resources, X.-J.W., G.B., B.R.; supervision, B.R.

## Declaration of generative AI and AI-assisted technologies in the writing process

Authors did not use generative AI or AI-assisted technologies in the writing process of this manuscript.

## Methods

### EXPERIMENTAL MODEL AND STUDY PARTICIPANT DETAILS

#### Mice

Female and male mice were used for all experiments (P30-P120) indiscriminately. The following primary transgenic mouse lines were used to generate the compound transgenic crosses described in the present study: Vip-ires-Flpo (Jax #028578) ^32^, Cck-ires-Cre (Jax #012706), Calb2-ires-Cre (Jax #010774; CR-Cre), Crh-ires-Cre (Jax #012704), Sncg-ires-Flpo, (Jax #034424), Ai65 (Cre + Flp dependent tdTomato; Jax #021875)^104^, Ai80 (Cre + Flp dependent CatCh-YFP; Jax #025109) ^105^, and RC:FLTG (Flp -> tdTomato; Flp + Cre -> EGFP; Jax #026932) ^38^, all obtained from the Jackson Laboratory. Additional lines included Lhx6(BAC)-EGFP (Tg(Lhx6-EGFP)BP221Gsat), ChAT-ires-Flpo (Jax #036281), and Ai80F (a Cre-deleted version of Ai80 made by us). The reporter lines Ai65, Ai80, Ai80F and FLTG were maintained as homozygous stocks before being crossed with compound driver lines to generate experimental animals. Genotyping was performed on a fee basis by the Genotyping Core Laboratory at the NYU Langone Medical Center. All experiments were conducted in accordance with protocols approved by the Division of Comparative Medicine at the NYU Grossman School of Medicine.

### METHOD DETAILS

#### Histology and Immunohistochemistry

To obtain brain tissue for histological analysis, animals (P40–P60) underwent transcardial perfusion with 4% paraformaldehyde (PFA) in PBS (diluted from a 32% stock; Electron Microscopy Sciences, cat#15714). Following perfusion, brains were dissected and subjected to a post-dissection fixation period of 0–16 hours. For thin cryosection preparation, brains were equilibrated overnight in 30% sucrose/PBS, then embedded in Tissue-Plus O.C.T. compound (Scigen, cat#4583) and frozen. Cryosections (20 μm) were cut using a Leica CM3050 cryostat, mounted onto glass slides (Shandon ColorFrost Plus; Thermo-Scientific, cat#9991013), and stored at –20°C until further processing. Thicker tissue sections (50-300 μm) were prepared from fixed brains in PBS using a vibratome (Leica). Some 300 μm-thick slices, initially prepared for electrophysiological recordings, were also used for histological analysis to confirm CR expression in patched neurons.

For anti-CR immunohistochemistry (IHC) on free floating vibratome brain sections, tissue sections were fixed for 15 min in 4% PFA in 1xPBS at room temperature, then washed in PBS, then blocked for 1 hour at room temperature in PBS containing 0.3% Triton X-100 and 2% normal donkey serum. Sections were incubated for two days at 4°C with the primary antibody (mouse anti-CR; EMD Millipore, MAB1568; 1:1500 dilution) and 0.2% streptavidin-AlexaFluor 647 in blocking solution. Following washes in PBS, sections were incubated for 2 hours at room temperature with the secondary antibody (donkey anti-mouse AlexaFluor 488; Invitrogen, cat#A-21202; 1:1000 dilution). Immunohistochemistry on thin cryosections was performed as described previously^49^, using rabbit anti-tdTomato (Rockland; cat# 600-401-379) and rat anti-EGFP (Nacalai USA; cat# 04404-84) primary antibodies to enhance tdTomato and EGFP signal, respectively (e.g., in FLTG reporter crosses).

#### Morphological reconstructions

Neurons were recorded using whole-cell electrophysiology while being held in an internal solution containing ∼0.4% or ∼1.0% biocytin for a minimum of 15 minutes. In cases where neurons were maintained for close to 15 minutes, slices were briefly returned to the incubation chamber to enhance biocytin diffusion throughout the recorded cell. After recordings, slices were fixed in 4% paraformaldehyde/PBS (prepared from a 32% stock; Electron Microscopy Sciences, cat#15714) and stored at 4°C for 1–7 days. Following fixation, slices were washed thoroughly in PBS and incubated overnight at 4°C in a 0.4% streptavidin-AlexaFluor 647 solution (Invitrogen; 498 µL 0.3% Triton X-100 in PBS and 2 µL streptavidin per slice). After incubation, slices were washed again in PBS, mounted on glass slides with Fluoromount-G (Invitrogen), and imaged using a Zeiss confocal microscope with a 40x–63x oil-immersion objective. Neuronal morphology was reconstructed in three dimensions using Neurolucida, enabling comprehensive analysis of cellular morphology.

#### Stereotaxic injections

Both male and female mice, aged P25–P45 at the time of injection, were used in these experiments. Mice were initially anesthetized with isoflurane (∼3%; 1 L/min O₂ flow for 2 minutes) before being head-fixed onto a stereotaxic frame (Kopf, Model 1900) using non-rupture ear bars. Anesthesia was maintained at 2% isoflurane throughout the procedure, and body temperature was regulated using an electronic warming pad. Under aseptic conditions, the skull was exposed, and a small craniotomy was performed over the barrel cortex (VS1) (coordinates: –0.90 mm AP from Bregma, ±3.20 mm ML, and 0.60 mm DV from pia) or vibrissae primary motor cortex (vM1) (coordinates: +0.80 mm AP from Bregma, ±0.25 mm ML, and 0.50 mm DV from pia) using a thin drill. Injections were delivered using a Nanoject III pressure injector (Drummond Scientific) mounted on the stereotaxic frame. A pulled borosilicate glass injection pipette (1B100-4; World Precision Instruments) with a ∼35 mm taper length and a ∼40 μm tip diameter was used for precise delivery of viral preparations at a rate of 4 nL/min, with injection volumes ranging from 50–200 nL.

#### Slice Preparation

Adult transgenic mice of either sex (postnatal day 30–120; mean age = 44 days) were terminally anesthetized with isoflurane. Once unresponsive, mice underwent transcardial perfusion with ice-cold sucrose-based artificial cerebrospinal fluid (sucrose-ACSF) containing (in mM): 87 NaCl, 75 sucrose, 2.5 KCl, 26 NaHCO₃, 1.25 NaH₂PO₄, 10 glucose, 1.0 CaCl₂, and 2.0 MgCl₂, continuously saturated with 95% O₂ / 5% CO₂. Following perfusion, mice were decapitated, and the brains were rapidly extracted. The caudal portion of the brain was affixed to a slicing stage, positioning the rostral end at a 15° forward pitch. The stage was then submerged in a chamber filled with ice-cold, oxygenated sucrose-ACSF, and 300 µm-thick coronal brain sections were obtained using a Leica VT1200S vibratome. Brain slices were then incubated at 35°C for 30 minutes in either the same sucrose-based solution or the recording-ACSF (described in the electrophysiology section). Slices were subsequently maintained at room temperature for at least 1 hour before the start of electrophysiological recordings.

#### Electrophysiological Recordings

Brain slices were transferred to a recording chamber continuously perfused with artificial cerebrospinal fluid (ACSF) containing (in mM): 120 NaCl, 2.5 KCl, 25 NaHCO₃, 1.4 NaH₂PO₄, 21 glucose, 0.4 Na-ascorbate, 2 Na-pyruvate, 2 CaCl₂, and 1 MgCl₂, saturated with 95% O₂ / 5% CO₂ and maintained at 29–32°C. In select experiments, the bath solution was supplemented with the NMDA receptor antagonist D-AP5 (25 µM; Abcam) and/or the AMPA receptor antagonist CNQX (10 µM; Abcam). Neurons were visualized using differential interference contrast (DIC) optics and fluorescence illumination from an LED power source (Mightex) on an upright Olympus microscope (BX50WI or BX51WI) to identify tdTomato and/or GFP-expressing neurons. All recordings were conducted in the barrel field of the primary somatosensory cortex (S1BF), specifically in layers 2–4. Cortical layers were identified as mentioned in morphological analysis methods. Whole-cell patch-clamp recordings were performed in current-clamp and voltage-clamp modes using an internal solution containing (in mM): 130 K-gluconate, 10 HEPES, 1.1 EGTA, 2 Mg-ATP, 0.4 Na-GTP, 10 Na-phosphocreatine, 1.5 MgCl₂, and 0.3–0.5% or 0.9–1.1% biocytin. The solution was adjusted to pH 7.3 using 1 M KOH. Recording pipettes were fabricated from borosilicate glass capillaries (inner/outer diameter: 1.5 mm / 0.86 mm) using a horizontal puller (Sutter Instruments), yielding pipette resistances of 2–6 MΩ. Before obtaining whole-cell access, a gigaseal was formed, and pipette capacitance was compensated. Access resistance was monitored throughout the experiment and fully compensated; neurons with access resistances >40 MΩ were excluded from intrinsic property analyses. Electrophysiological data were acquired using a MultiClamp 700B amplifier (Molecular Devices), a Digidata digitizer (1440A or 1550B series, Molecular Devices), and Clampex software (v10.6 or v10.7, Molecular Devices). Data were sampled at 20 kHz and low-pass filtered at 10 kHz.

#### Electrophysiological characterization

Neurons were recorded in the indicated mouse lines to characterize each VIP IN subpopulation. In addition, we recorded from L2/3 pyramidal cells (PCs) as well as from PV and SST INs to train an unbiased classifier to distinguish between these two types of INs electrophysiologically when characterizing the efferent connectivity of VIP IN subpopulations (see quantification and statistical analysis – PV and SST classifier). Neurons were characterized for intrinsic and active properties in current-clamp mode from a resting potential of ∼–70 mV. Neurons were injected with 1 s long square pulses of increasing current.

#### Optogenetics in acute brain slices and DSI

For optogenetic experiments, layer 2/3 PCs were first identified visually using DIC microscopy, then patched in whole-cell mode and confirmed as PCs through brief electrophysiological characterization. Blue light (470 nm) pulses (2 ms, >100 mA) were delivered to the slice using TTL-triggered pulses directed through a Mightex LED system under 40× magnification, with the PC soma centered in the field of view. Optically evoked inhibitory postsynaptic currents (IPSCs) were recorded under voltage-clamp mode at a holding potential of –50 mV, with D-AP5 (25 μM) and CNQX (10 μM) included in the bath solution to block glutamatergic transmission. If a connection was detected, the DSI protocol was executed as described in Figure 5.

#### Silicon probe implantation and recordings

Mice (n=5, 28–35 g, 3–10 months old) were implanted with 64-site silicon probes (NeuroNexus, Cambridge NeuroTech or Diagnostic Biochips) in the PPC (AP 2.0 mm, ML 1.75 mm, DL 0.6 mm), as described previously ^106^. Ground and reference wires were implanted in the skull above the cerebellum, and a grounded copper mesh hat was constructed, protecting, and electrically shielding, the probes. Probes were mounted on plastic microdrives that were advanced to layer 6 over the course of 5–8 d after surgery. A 100–200 μm fiber optic was attached to one of the shanks of the silicon probe. After implantation, animals were allowed to recover for at least 1 week and were housed individually under standard conditions (71–73 °F/21.5-23 °C and 40–50% relative humidity) in the animal facility and kept on a 12 h reverse light/dark cycle. We recorded the mice while they slept or walked around freely in the home cage, and the recording session started 1–2 hr after the onset of the dark phase.

### QUANTIFICATION AND STATISTICAL ANALYSIS

#### Morphological analysis

Neurolucida Explorer was used to perform Sholl analysis on reconstructed neurons (n = 32) and to extract morphological parameters, including somatic surface area and volume, number of primary dendrites, total dendritic length, mean dendritic length, dendritic surface area, volume occupied by dendrites, dendritic polarity, total axonal length, axonal surface area, volume occupied by the axon, and axonal polarity. These parameters were subsequently analyzed using a custom MATLAB routine. Somatic localization within cortical layers was determined by identifying either cortical columnar boundaries or layer borders, followed by quantification of the normalized position within the established limits. Layers were visually identified in confocal images based on anatomical features: (A) The L1-L2 border was defined by a sharp increase in soma density from L1. (B) The L3-L4 border was identified by the top of the barrels and the absence of pyramidal cells (PCs) characteristic of L2/3. (C) L5a appeared as a distinct band below L4. (D) The L5b-L6 border was delineated by a sharp drop in pyramidal cell density. Unbiased morphological clustering was performed using Ward’s method based on all extracted morphological parameters. The Gap Statistic was used to determine the optimal number of clusters, yielding values between k=2 and k=3 across iterations. Given the stability of the clustering structure and biological interpretability, we selected k=3 with a distance threshold of 32 for final group separation. Axonal and dendritic layer distributions were quantified by dividing each vectorized reconstructed neuron into layers L1–L6. Next, vectors were isolated per layer and binarized using FIJI/ImageJ, generating a layer-dependent profile for each neuron. These profiles were further analyzed using a custom MATLAB routine.

#### PV & SST IN Classifier

To determine the output connectivity of VIP IN subpopulations to PV and SST INs, an additional reporter allele, Lhx6(BAC)-EGFP, was utilized to label these two IN populations. In order to differentiate between the two types of INs, we used an unbiased classifier that was trained on a set of electrophysiological data from 291 identified INs (125 PV INs and 166 SST INs). The reference cell datasets were from this laboratory as well as from the Allen Institute ^33^. The datasets from this laboratory were all from INs in L2/3 of vS1. The viral and genetic methods used to label reference INs included: PV-Cre;Ai9, PV-Cre + injection of Cre-dependent AAV bearing a fluorophore reporter, the G42 and B13 GFP mouse lines, SST-Cre;Ai9, SST-Flpo + injection of Flp-dependent AAV bearing a fluorophore reporter, SST-Cre + injection of Cre-dependent AAV bearing a fluorophore, injection of AAV-S5E2-dTom-nlsdTom (AAV PHP.eB) virus to label PV INs ^107^, injection of AAV-S9E10-dTom virus to label SST INs ^108^, or injection of AAV9-Sst44-nls-mScarlet virus to label SST INs ^91,109^. Reference cell data from the Allen Institute ^35^ of L2/3 of the mouse primary visual cortex provided cells that were transcriptomically identified as PV or SST INs. We trained a bagging classifier model on the following six electrophysiological parameters from the reference cells: adaptation index, firing rate, membrane time constant, input resistance, rheobase, and AP half-width. Using 5-fold cross-validation, the model had an accuracy of 94.2%. Additionally, we verified the validity of combining the IN datasets by fitting the same bagging classifier model to the Allen Institute dataset alone and predicting our lab’s dataset. The model achieved similar accuracy on both datasets (>91%).

#### Unit activity analysis

Spike sorting was performed semi-automatically with KiloSort 1 (https://github.com/cortex-lab/KiloSort; RRID:SCR_016422), as previously described ^106^ and using our own pipeline KilosortWrapper (a wrapper for KiloSort, DOI;; ^110^). This was followed by manual adjustment of the waveform clusters using the software Phy 2 (https://github.com/kwikteam/phy; ^111^) and plugins for phy designed in the laboratory (https://github.com/petersenpeter/phy-plugins; ^112^). Unit clustering generated two clearly separable groups based on their spike autocorrelograms, waveform characteristics and firing rate ^106^. Pyramidal cell (PC), and interneurons (INs) were tentatively separated based on these two clusters. A more reliable cell identity was assigned after inspection of all features, assisted by monosynaptic excitatory and inhibitory interactions between simultaneously recorded, well-isolated units and light responses ^106,113^. Units were defined as optogenetically modulated cells based on their significant modulation against randomly shuffled pulse times (500 replicates) and testing for significant difference between the observed value and the random distribution.

#### Statistical analysis

Statistical analyses were performed using standard MATLAB functions and GraphPad Prism. Sample size was not determined by a formal power analysis; however, the number of animals, trials, and *in vitro* and *in vivo* recorded cells were comparable to or exceeded those used in previous studies ^113^. All data were obtained from experimental replicates, with a minimum of four independent experimental repeats per assay. Replication attempts were consistently successful. Data collection was not conducted under blinded conditions regarding subject groups, but data analysis was performed either blinded to the scorer or in cases where manual scoring was unnecessary. The Lilliefors test was applied to assess normality, guiding the selection of appropriate statistical tests. For normally distributed data, statistical comparisons were conducted using unpaired or paired Student’s t-tests for two-group comparisons and one-way analysis of variance (ANOVA) followed by Bonferroni’s post hoc test for multiple group comparisons. For non-normally distributed data or when assumptions of parametric tests were not met, statistical analyses were performed using non-parametric tests, including the two-tailed Wilcoxon paired signed-rank test for paired data, the Mann-Whitney U test for unpaired comparisons, and the Kruskal-Wallis one-way analysis of variance for multiple group comparisons. For post hoc multiple comparisons, the Dunn-Sidak correction was applied. Statistical significance was defined as *p* < 0.05, with additional thresholds indicated as p < 0.01, p < 0.001, and p < 0.0001. For box-and-whisker plots the box represents the interquartile range (IQR: 25th to 75th percentile), the horizontal line within the box indicates the median, and whiskers extend to 1.5x the IQR unless outliers are present.

### KEY RESOURCES TABLE

| REAGENT or RESOURCE | SOURCE | IDENTIFIER |
| --- | --- | --- |
| Antibodies |  |  |
| Mouse anti-Calretinin (clone 6B8.2) | EMD Millipore | Cat# MAB1568;<br>RRID:AB_94259 |
| Rabbit anti-tdTomato | Rockland Immunochemicals | Cat# 600-401-379;<br>RRID:AB_2209751 |
| Rat anti-EGFP (clone GF090R) | Nacalai USA | Cat# 04404-84;<br>RRID:AB_10013361 |
| Donkey anti-Mouse IgG (H+L), Alexa Fluor 488 | Invitrogen | Cat# A-21202;<br>RRID:AB_141607 |
| Streptavidin, Alexa Fluor 647 Conjugate | Thermo Fisher Scientific | Cat# S32357;<br>RRID:AB_2336066 |
| Recombinant DNA (Viral Constructs) |  |  |
| AAV-hSyn-hChR2(H134R)-EYFP | Addgene plasmid #26973 | Addgene_26973 |
| AAV pCAG-FLEX-tdTomato-WPRE | Addgene plasmid #51503 | Addgene_51503 |
| AAV-PHP.eB-S5E2-dTom-nls-dTom | Addgene plasmid #135630<sup>107</sup> | Addgene_135630 |
| AAV-S9E10-dTom | Fishell Lab<sup>108</sup> | N/A |
| AAV9-Sst44-nls-mScarlet | Harvey Lab<sup>91,109</sup> | N/A |
| Chemicals, peptides, and recombinant proteins |  |  |
| Paraformaldehyde (32% stock) | Electron Microscopy Sciences | Cat# 15714 |
| Tissue-Plus O.C.T. compound | Scigen US | Cat# 4583 |
| Biocytin | Sigma-Aldrich | Cat# B4261 |
| D-AP5 (NMDA receptor antagonist) | Abcam | Cat# ab120003 |
| CNQX (AMPA receptor antagonist) | Abcam | Cat# ab120017 |
| SR95531 (Gabazine) | Tocris Bioscience | Cat# 1262 |
| Noradrenaline bitartrate | Tocris Bioscience | Cat# 5169 |
| Triton X-100 | Sigma-Aldrich | Cat# T9284 |
| Normal donkey serum | Jackson ImmunoResearch | Cat# 017-000-121 |
| Fluoromount-G | Invitrogen | Cat# 00-4958 |
| Experimental models: Organisms/strains |  |  |
| Mouse: Vip-ires-Flpo | The Jackson Laboratory | JAX:028578;<br>RRID:IMSR_JAX:028578 |
| Mouse: Cck-ires-Cre | The Jackson Laboratory | JAX:012706;<br>RRID:IMSR_JAX:012706 |
| Mouse: Calb2-ires-Cre | The Jackson Laboratory | JAX:010774;<br>RRID:IMSR_JAX:010774 |
| Mouse: Crh-ires-Cre | The Jackson Laboratory | JAX:012704;<br>RRID:IMSR_JAX:012704 |
| Mouse: Sncg-ires-Flpo | The Jackson Laboratory | JAX:034424;<br>RRID:IMSR_JAX:034424 |
| Mouse: Ai65 (tdTomato reporter) | The Jackson Laboratory | JAX:021875;<br>RRID:IMSR_JAX:021875 |
| Mouse: Ai80 (CatCh-YFP reporter) | The Jackson Laboratory | JAX:025109;<br>RRID:IMSR_JAX:025109 |
| Mouse: RC:FLTG (Flp→tdTomato; Flp+Cre→EGFP) | The Jackson Laboratory | JAX:026932;<br>RRID:IMSR_JAX:026932 |
| Mouse: Lhx6(BAC)-EGFP | GENSAT / MMRRC | RRID:MMRRC_000246-MU |
| Mouse: ChAT-ires-Flpo | The Jackson Laboratory | JAX:036281;<br>RRID:IMSR_JAX:036281 |
| Mouse: Ai80F (Cre-deleted Ai80) | This paper (in-house) | N/A |
| Software and algorithms |  |  |
| ImageJ | https://imagej.nih.gov/ij/<sup>114</sup> | RRID:SCR_003070 |
| Phy 2.0 b6 | GitHub;<br>https://github.com/kwik-team/phy | N/A |
| KiloSort | cortex-lab GitHub;<br>https://github.com/cortex-lab/KiloSort | RRID:SCR_016422 |
| KilosortWrapper | brendonw1 GitHub;<br>https://github.com/brendonw1/KilosortWrapper | N/A |
| Clampex (pClamp v10.6/10.7) | Molecular Devices | RRID:SCR_011323 |
| MATLAB | The MathWorks;<br>https://www.math-works.com/ | RRID:SCR_001622 |
| GraphPad Prism | GraphPad Software;<br>https://www.graphpad.com/ | RRID:SCR_002798 |
| Other |  |  |
| 64-site silicon probes (A4x16-5mm-50-703) | NeuroNexus / Cambridge NeuroTech / Diagnostic Bi-ochips | A4x16-5mm-50-703 |
| Fiber optic (200 μm) | Thorlabs | BFH48-200 |

