## Supplemental Material for "Inhibitory and disinhibitory VIP IN-mediated circuits in neocortex"

#### Supplemental Figure 1

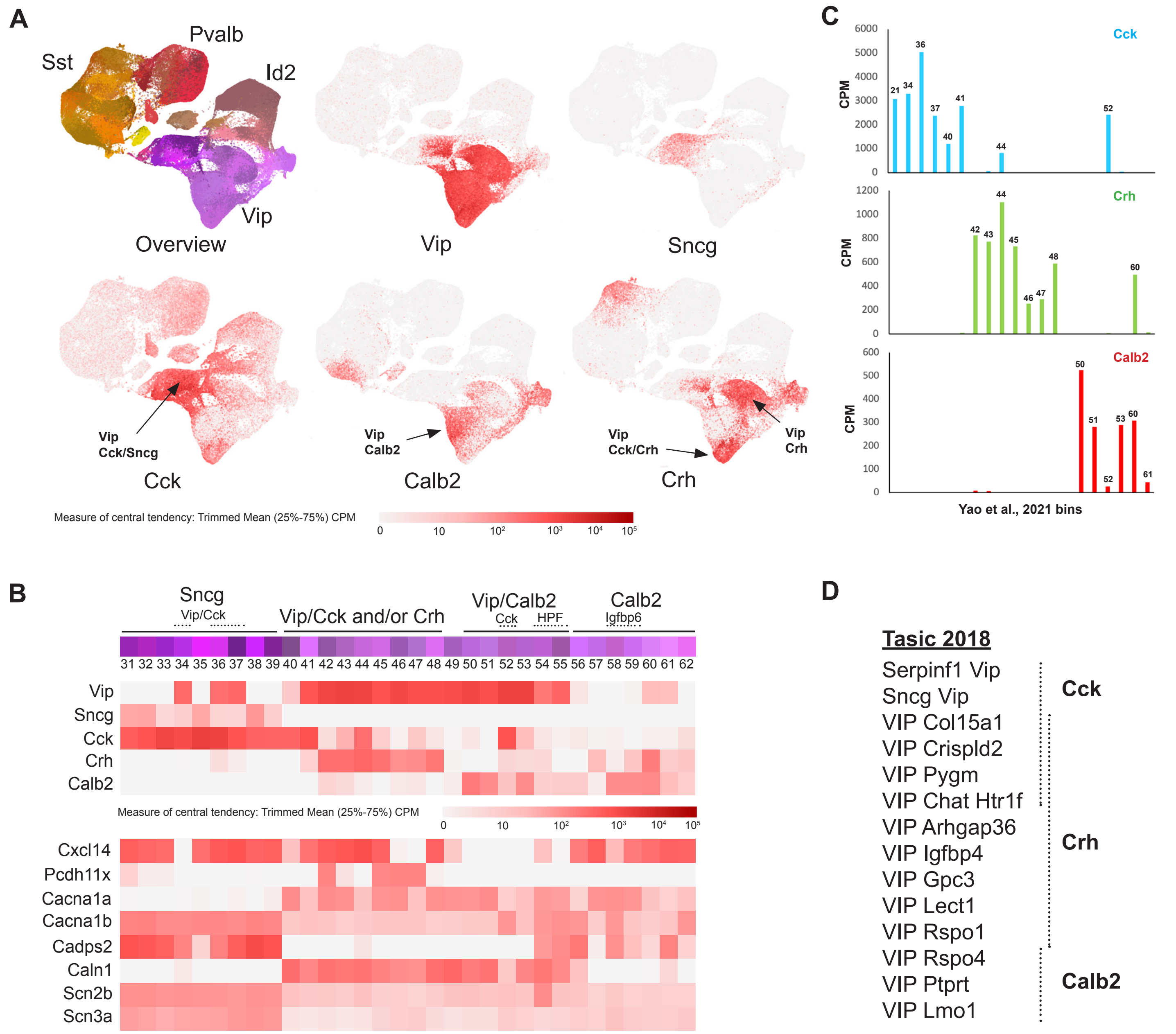

**Supplementary Figure 1. VIP IN transcriptomics.**

**(A)** Scatterplots of GABAergic INs from neocortex and hippocampus (Allen Institute scRNAseq data; Yao et al., 2021[1]), with overview and plots of Vip, Sncg, Cck, Calb2 (CR) and Crh expression (red color intensity illustrates trimmed mean 25-75% counts per million of each transcript; see color scale bar for conversion from log2 values). **(B)** Heat maps of Vip, Sncg, Cck, Crh, and Calb2 (CR) trimmed mean (25-75%) expression across the Yao et al., 2021[1] clusters, along with annotation (adapted from Machold and Rudy, 2024). Additional gene profiles are shown for Cxcl14, Pcdh11x (notably enriched in VIP/non-CCK non-CR), calcium channel subunits Cacna1a (non-VIP/Sncg) and Cacna1b (VIP/Sncg), calcium-related genes Cadps2 (VIP/Sncg) and Caln1 (non-VIP/Sncg), and sodium channel subunits Scn2b and Scn3a (VIP/Sncg). **(C)** Graphs of Cck, Crh and Calb2 (CR) mRNA expression (trimmed mean counts per million; CPM) across the Yao et al., 2021 [1] clusters. **(D)** Approximate correspondence between the VIP IN subtype categories described in Tasic et al., 2018 [2] and VIP/CCK, VIP/CRH, and VIP/CR subtypes.

### Supplemental Figure 2

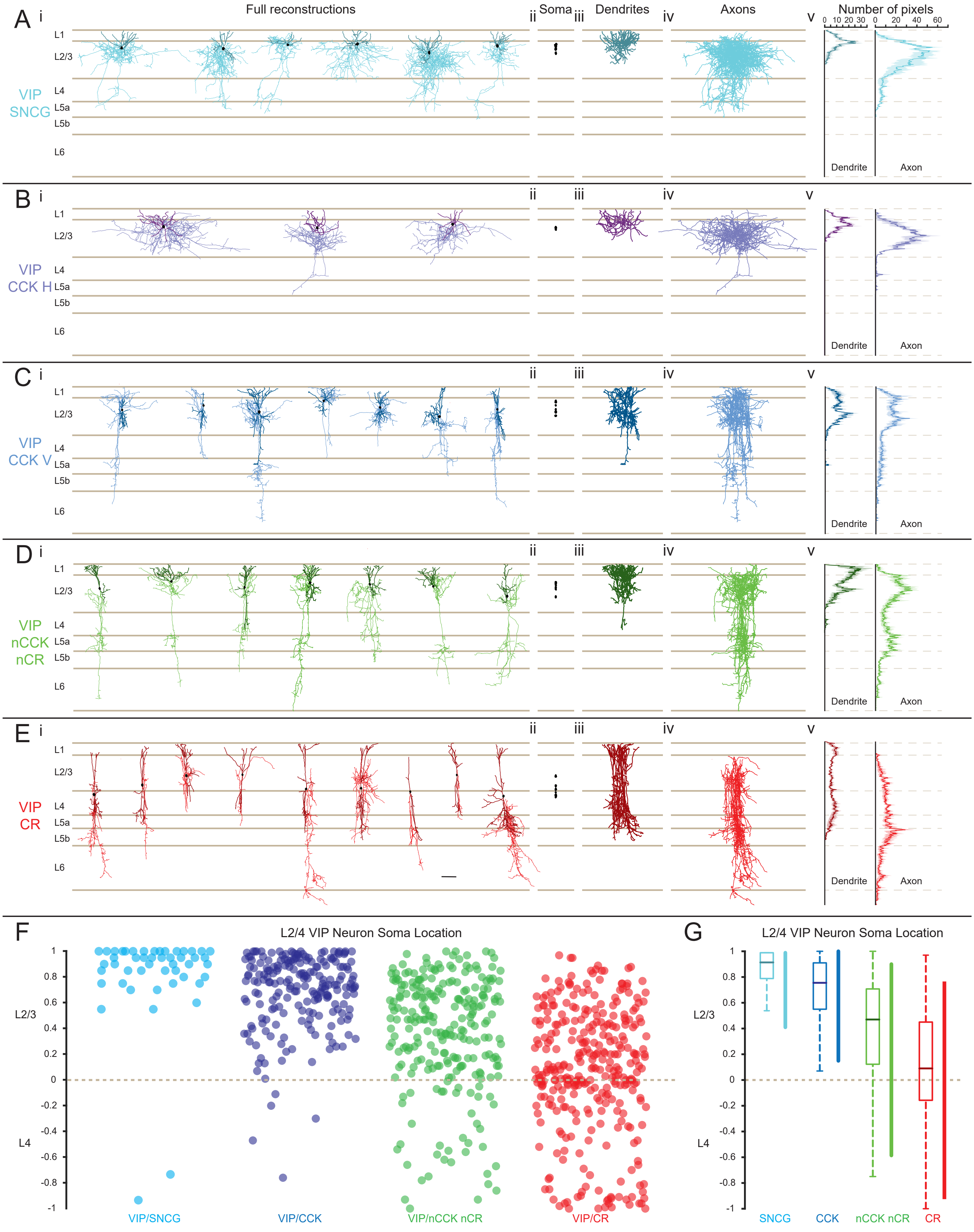

**Supplementary Figure 2. Morphological Reconstructions of VIP INs.**

Neurolucida reconstructions of all biocytin-filled neurons**. (A)** Reconstructions of SNCG-expressing VIP neurons (n = 6) from VIP-Cre/SNCG-Flp mice, showing the distribution of dendrites (dark cyan), axons (light cyan), and soma (black) across cortical layers **(i).** Superimposed somata of all reconstructed neurons **(ii)**. Superimposed dendrites of all reconstructed neurons **(iii)**. Superimposed axons of all reconstructed neurons **(iv)**. Quantification of pixel distribution across the cortical column for total reconstructed dendrites (left) and axons (right) **(v)**. **(B)** Same as (A) for CCK-expressing horizontal (CCK H) VIP neurons (n = 3) from VIP-Flp/CCK-Cre/FLTG mice with dendrites shown in dark lavender, axons shown in light lavender, and soma shown in black grouped with VIP/SNCG. **(C)** Same as (A) for CCK-expressing vertical (CCK V) VIP neurons (n = 7) reconstructed from VIP-Flp/CCK-Cre/FLTG mice with dendrites shown in dark blue, axons shown in light blue, and soma in black. **(D)** Same as (A) for non-CCK, non-CR neurons VIP neurons (n=7) from VIP-Flp/CCKCR-Cre/FLTG mice with dendrites shown in dark green,  axons shown in light green, and soma in black. **(E)** Same as (A) for CR-expressing VIP INs (n = 9) from VIP-Flp/CR-Cre/FLTG mice with dendrites shown in dark red, axons shown in light red, and soma in black. **(F)** Normalized soma location across layer 2/3 (1 to 0) and layer 4 (0 to -1) of VIP/SNCG (cyan), VIP/CCK (blue), VIP/nCCK nCR (green), and VIP/CR (red) (n=6-10 slices from 3-5 animals). **(G)** Summary plot of normalized soma location of VIP populations. Vertical bar at the right of box and whisker plot determines where 90% of the somas are located in layer 2/4.

### Supplemental Figure 3

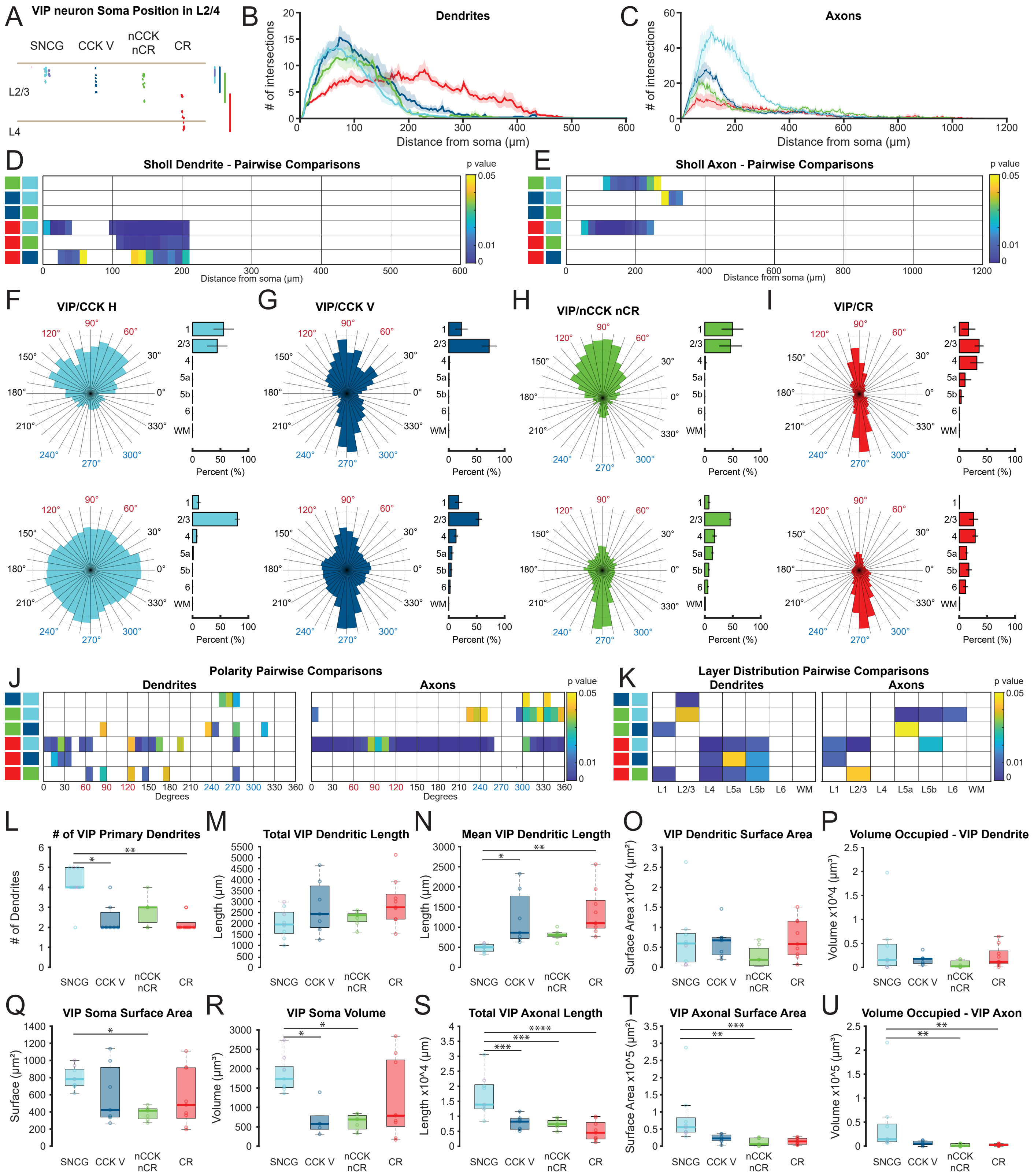

**Supplementary Figure 3. Quantification of morphological parameters of VIP IN populations.**

**(A)** Soma position of reconstructed cells VIP/SNCG (cyan) (VIP/CCK H indicated in lavender), VIP/CCK V (dark blue), VIP/nCCK nCR (green), and VIP/CR (red) morphologically reconstructed neurons throughout layer 2/4. **(B)** Dendritic Sholl analysis of VIP IN populations. Shaded area delineates SEM. **(C)** Axonal Sholl analysis of VIP IN populations. Shaded area delineates SEM. **(D)** Pairwise comparisons for the dendritic sholl analysis (Kruskal-Wallis test; p= 2.0187e-10; post hoc Dunn-Sidak). Colored squares at the left indicate the pairwise comparison of cells (VIP/SNCG (cyan), VIP/CCK H (lavender), VIP/CCK V (dark blue), VIP/ nCCKnCR (green), and VIP/CR (red). **(E)** Same as in **D** for the axonal sholl analysis (Kruskal-Wallis test; p= 0.0054215; post hoc Dunn-Sidak). **(F)** (Left; top) Mean dendritic polarity of VIP/CCK H showing their directional bias within the cortical column. In this representation, 90° corresponds to dendritic orientation toward the pia, whereas 270° corresponds to orientation toward the white matter. (Left; bottom) Same as in top but for axons. (Right; top) Layer-wise percent distribution of VIP/SNCG dendrites. (Right; bottom) Same as in top but for axons. Error bars represent SD. **(G)** Same as F but for VIP/CCK V. **(H)** Same as F but for VIP/nCCK nCR. **(I)** Same as F but for VIP/CR. **(J)** Pairwise comparisons for dendritic (left) and axonal (right) polarity (Kruskal-Wallis test; $p=1.7327163378x10^{-16}$; post hoc Dunn-Sidak). Colored squares at the left indicate the pairwise comparison of cells (VIP/SNCG (cyan), VIP/CCK V (dark blue), VIP/nCCKnCR (green), and VIP/CR (red) neurons). **(K)** Same as in **I** for dendritic (left) and axonal (right) layer distributions (Kruskal-Wallis test; dendrites $p= 1.7057686680x10^{-2}$ and axons $p=4.5067965911x10^{-3}$; post hoc Dunn-Sidak). (**L)** Number of Primary Dendrites (VIP/SNCG n=9; VIP/CCK V n=7; VIP/nCCKnCR n=7; VIP/CR H n=9). VIP/SNCG (VIP/CCK H indicated in lavender circles in VIP/SNCG group) neurons exhibited significantly more primary dendrites compared to VIP/CCK V and VIP/CR neurons (data not normally distributed, Kruskal-Wallis test, p= 0.0012316; Dunn-Sidak post hoc test, VIP/SNCG vs. VIP/CCK V p= 0.013372; VIP/SNCG vs. VIP/CR p= 0.0014117). **(M)** Total Dendritic Length. Total dendritic length remained similar across groups (data normally distributed, ANOVA, p=0. 12716). **(N)** Mean Dendritic Length. VIP/SNCG neurons exhibited shorter dendritic branches than other VIP INs (data normally distributed, ANOVA, p= 0.001271; Tukey-Kramer post hoc test, VIP/SNCG vs. VIP/CCK V p= 0.016074; VIP/SNCG vs. VIP/CR p= 0.0013568). **(O)** Dendritic Surface Area. Differences in dendritic surface area approached significance (data normally distributed, ANOVA, p= 0.31534). **(P)** Dendritic Volume. Dendritic volume remained consistent across VIP INs (data not normally distributed, Kruskal-Wallis, p= 0.12118). **(Q)** Soma Surface Area. Soma surface area did significantly differ among subtypes (data normally distributed, ANOVA, p= 0.028147; Tukey-Kramer post hoc test, VIP/SNCG vs. VIP/nCCK nCR p= 0.016362). **(R)** Soma Volume. Soma volume was significantly different across groups (data not normally distributed, Kruskal-Wallis test, p= 0.0032016; post hoc Dunn-Sidak, VIP/SNCG vs. VIP/nCCK nCR p= 0.014069, VIP/SNCG vs. VIP/CCK V p= 0.014275). **(S)** Total Axonal Length. VIP/SNCG neurons exhibited significantly longer total axonal length compared to other subtypes (data normally distributed, ANOVA, $p= 9.9012x10^{-6}$; Tukey-Kramer post hoc test, VIP/SNCG vs. VIP/CCK V p= 0.00090171, VIP/SNCG vs. VIP/nCCKnCR p= 0.00053172, VIP/SNCG vs. VIP/CR $p= 9.792x10^{-6}$). **(T)** Axonal Surface Area. Axonal surface area was significantly greater in VIP/SNCG neurons (data not normally distributed,

Kruskal-Wallis, p= 0.00021431; post hoc Dunn-Sidak, VIP/SNCG vs. VIP/CCK V reaching significance p= 0.057511, VIP/SNCG vs. VIP/nCCKnCR p= 0.001147, VIP/SNCG vs. VIP/CR p= 0.00080323). **(U)** Axonal Volume. VIP/SNCG neurons had a significantly greater axonal volume compared to other groups (data not normally distributed, Kruskal-Wallis, p= 0.00037299; post hoc Dunn-Sidak, VIP/SNCG vs. VIP/nCCKnCR p= 0.0013626, VIP/SNCG vs. VIP/CR p= 0.0016141). These results suggest that while soma-related properties remain consistent across VIP IN populations, VIP/SNCG neurons are morphologically distinct with longer dendrites and significantly larger axonal length, surface area, and volume compared to other VIP IN populations.

**Supplemental Figure 4**

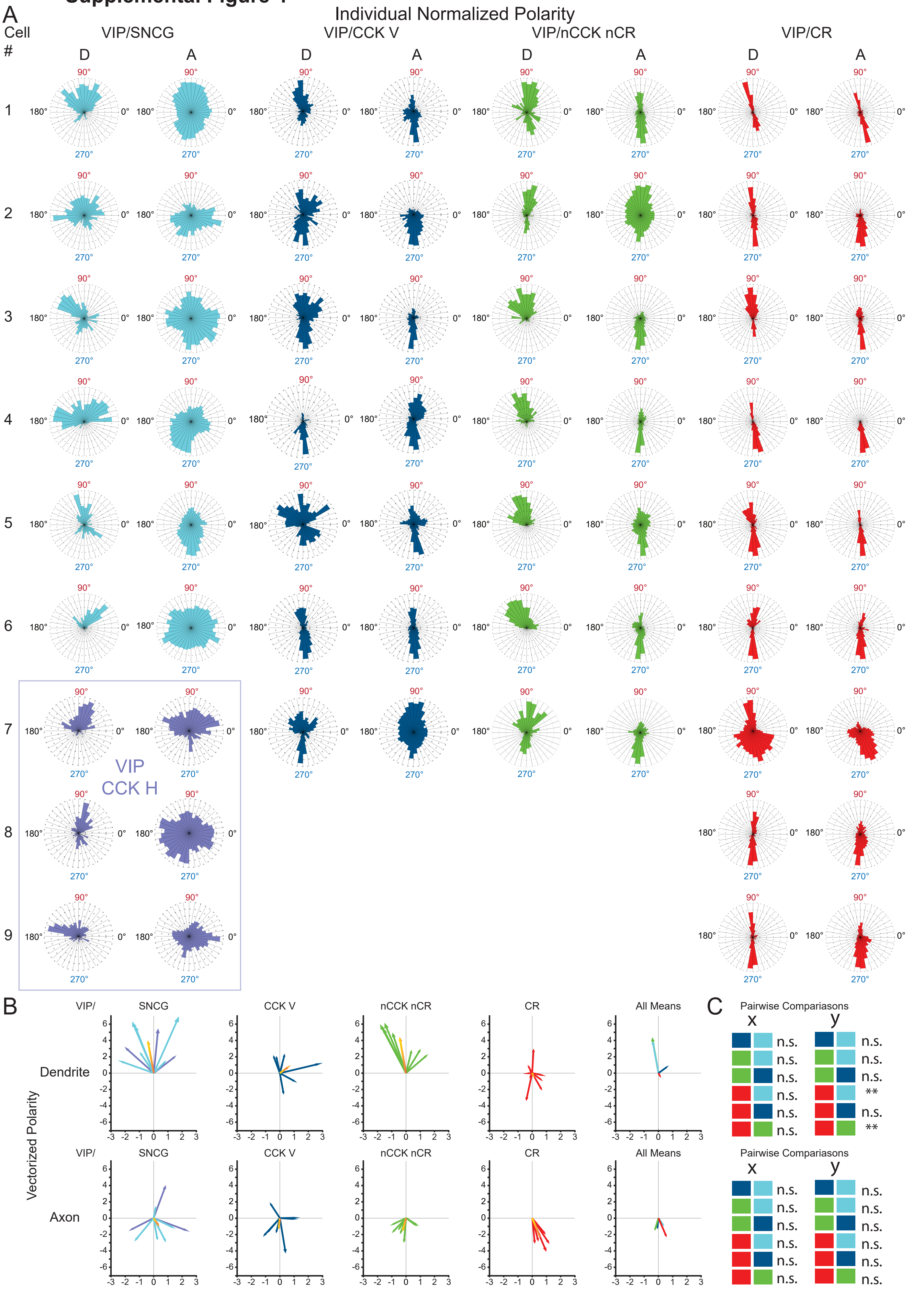

**Supplementary Figure 4. Individual Neurite Polarity of VIP IN populations.**

**(A)** Individual VIP cells polarity for dendrites (D) and axons (A) of VIP/SNCG (cyan), VIP/CCK H (lavender), VIP/CCK V (dark blue), VIP/nCCK nCR (green), and VIP/CR (red). **(B)** Vectorized dendritic and axonal polarity analysis. Vectorized dendritic (top) and axonal (bottom) polarity of individual neurons within each VIP population, with the mean vectorized polarity indicated in yellow-red. Summary of all VIP populations dendritic (last;top) and axonal (last;bottom) vectorized means. **(C)** Pairwise comparisons of vectorized dendritic (top) and axonal (bottom) polarities. The Kruskal-Wallis test was performed followed by a Dunn-Sidak post hoc test to compare the dendritic and axonal distributions among VIP/SNCG, VIP/CCK V, VIP/nCCKnCR, and VIP/CR neurons (Kurskal-Wallis - dendrites (x) $p = 0.56169$; dendrites (y) $p = 0.00035$; axons (x) $p = 0.23931$; axons (y) $p = 0.30261$). Dendrites of VIP/CR vs. VIP/SNCG and VIP/CR vs VIP/nCCK nCR were significantly different in only (y) components for dendrites (VIP/CR vs. VIP/SNCG $p = 0.00545$; VIP/CR vs VIP/nCCK nCR $p = 0.00273$).

#### Supplemental Figure 5

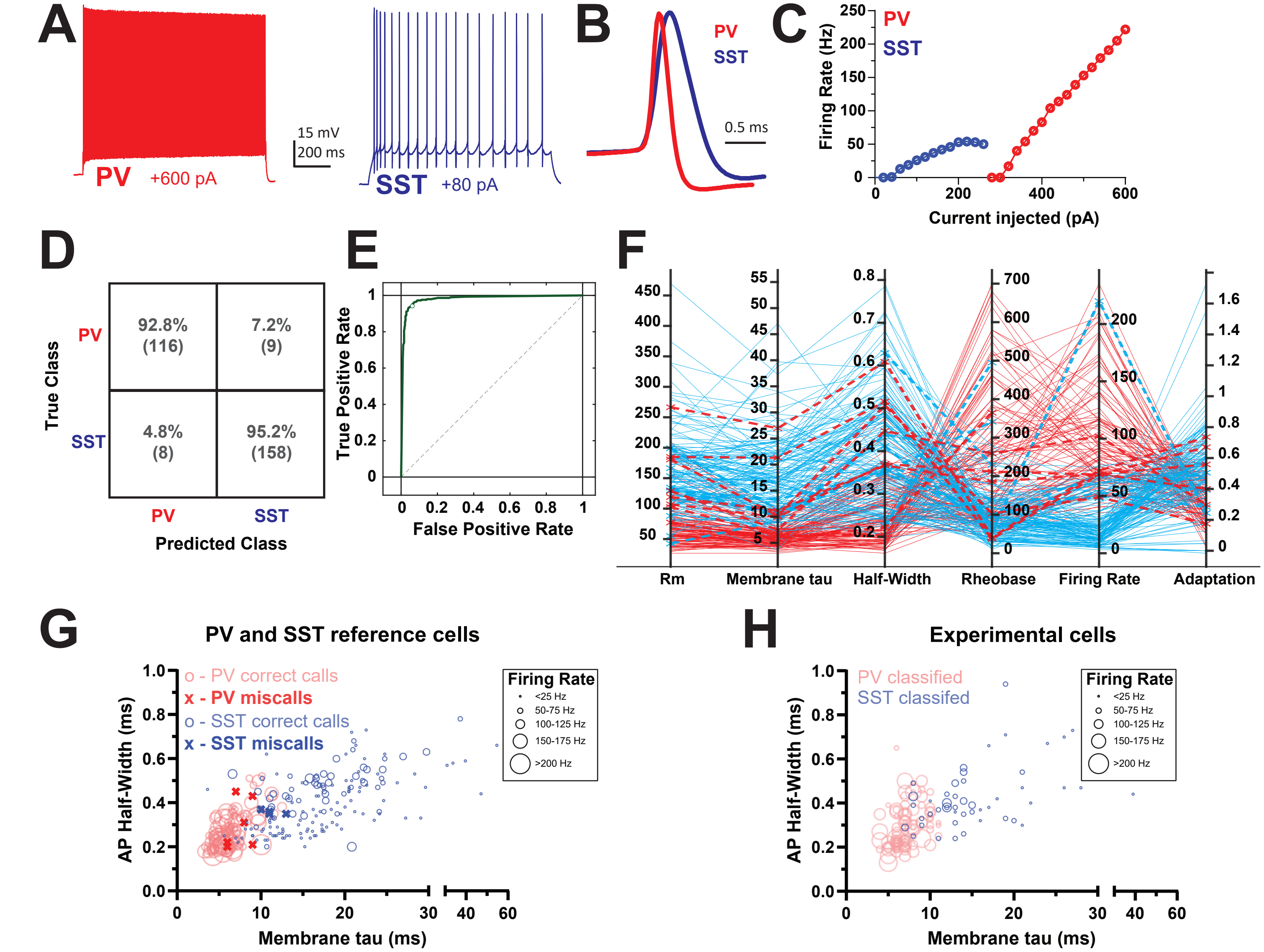

**Supplementary Figure 5. Differentiation of PV and SST INs in Lhx6 mice.**

**(A)** Example traces of firing patterns of a PV IN (red) and an SST IN (blue) showing their distinctive firing patterns, both at 2 x rheobase current levels. PV INs have a high firing rate and SST INs exhibit strong spike-frequency adaptation. The INs were fluorescently tagged cells in PV-Cre-Ai9 and SST-Cre-Ai9 animals. **(B)** Overlay of averaged and peak-normalized action potentials (APs) from the same cells as in **A** showing that APs from PV INs are narrower than those from SST INs. **(C)** Input-output firing rate curves for the same cells as in **A** showing that PV INs have higher rheobase and higher maximal firing rates than SST INs. **(D)** Confusion matrix for validating the behavior of an ensemble classifier on ground truth PV and SST INs. See methods for details on the classifier used. The classifier had an accuracy of 94.2%. **(E)** ROC curve for the ensemble classifier shown in **D**. **(F)** Parallel coordinates plot for several electrophysiological parameters for all reference PV (red) and SST (blue) INs on which the classifier was tested. Correctly identified INs indicated with solid lines and incorrectly identified INs are indicated by dashed lines. **(G)** Scatter plot of all reference PV and SST INs for three electrophysiological parameters: AP Half-Width, Membrane time constant (tau), and firing rate at 2 x rheobase, with correct and miscalls indicated by open circles or x's, respectively. **(H)** As in **G** but for Lhx6$^{+}$ INs used for VIP output connectivity (see Figure 4) and classified as PV or SST INs by the unbiased classifier (see Methods for details).
